# Sharp and smooth state transitions during resting state brain dynamics

**DOI:** 10.64898/2025.12.09.693219

**Authors:** Luiz Pessoa, Xiaoyu Zhou

## Abstract

Resting-state fMRI is routinely described in terms of a small set of recurring whole-brain states. Far less is known about what happens *between* them. Two pictures compete. In one, the brain *toggles*: it switches between qualitatively distinct regimes, and a transition is a discrete event. In the other, apparent states are convenient labels applied to a continuous trajectory, and transitions are mostly boundaries imposed by the analyst rather than events in the brain. We adjudicated between them with a multi-level Switching Linear Dynamical System fit to Human Connectome Project data (*N* = 500; 254 cortical and subcortical regions), in which every participant has their own equations of motion and switching parameters, tied to group-level parameters and estimated jointly with them. The model estimates the dynamics on either side of a boundary, so transitions can be characterized rather than merely counted. Transitions were structured and strongly heterogeneous. Four of the eight transitions examined reversed the estimated flow between two samples acquired 720 ms apart, two of them deeply, while others merely reoriented and were “one-sided”, with activity flat before the switch and changing only afterwards. Sharp and smooth transitions shared destination states and engaged overlapping networks, so the heterogeneity cannot be attributed simply to hemodynamic filtering. What predicted sharpness was the direction of travel, such that the brain drifts out of a dominant, long-dwell state regime and is sharply switched into it. To link these system-level dynamics to anatomy we introduce *transition importance*, which asks how much the model’s evidence for a particular switch depends on a region’s signal. The evidence accumulated over seconds before a switch, raising its odds by 1.8 to 3.2 times, and discharged immediately after. In several transitions the regions that steer a state shared no region with those that end it, so a region’s activity magnitude does not determine its contribution to system-level dynamics. Our findings suggest that spontaneous brain activity is organized as much by how the brain moves between configurations as by which ones it occupies, making transitions themselves an important target of study.

## Introduction

Functional MRI acquired in the absence of tasks is now commonly described in terms of a limited set of recurring, whole-brain patterns, or *states* (Allen et al., 2012; Vidaurre et al., 2017; Greene et al., 2023). The claim is supported by methods that differ considerably in their assumptions, including sliding-window correlation plus clustering (Allen et al., 2012; Hutchison et al., 2013), Hidden Markov Models (Vidaurre et al., 2017, 2018a; Song et al., 2023), and co-activation pattern analyses (Liu et al., 2018; Karahanoğlu and Van De Ville, 2015; Peng et al., 2023; Gutierrez-Barragan et al., 2024; Lee et al., 2024). Although the specific state repertoires reported vary, the agreement across techniques is substantial, and the field has converged on a vocabulary of states, dwell times, occupancies, and transition probabilities.

Two rather different pictures of resting-state dynamics are compatible with the existing evidence. The first picture is that the brain *toggles*. On this view, large-scale activity settles into one of several quasi-stable configurations and leaves it abruptly, so that the change of configuration is fast relative to the time spent in either. This is the picture suggested by theoretical work on metastability, itinerancy among attractors, and heteroclinic dynamics (Rabinovich et al., 2006; Tognoli and Kelso, 2014; Miller, 2016; Deco and Jirsa, 2012; Cabral et al., 2014), and it has direct empirical precedent at finer spatial and temporal scales. Ensembles in sensory cortex progress through sequences of states whose boundaries are abrupt when examined on single trials but are smeared by averaging (Jones et al., 2007; Mazzucato et al., 2015; La Camera et al., 2019; Recanatesi et al., 2022). Magnetoencephalography analyzed with Hidden Markov Models reveals network states lasting 50–200 ms with fast onsets and offsets (Baker et al., 2014; Vidaurre et al., 2018b). And the nervous system demonstrably implements genuine toggles: cortical up and down states (Sanchez-Vives and McCormick, 2000), the mutually inhibitory flip-flop circuits that produce sharp sleep–wake transitions (Saper et al., 2010), and the nonlinear “ignition” posited by global workspace accounts of conscious access (Dehaene and Changeux, 2011).

The second picture is that resting-state dynamics is essentially continuous, and that states are a discretization of a smooth trajectory. Quasi-periodic patterns propagate across cortex over 10 to 30 seconds, carrying activity gradually from default-mode-like to task-positive-like configurations and back (Majeed et al., 2011; Keilholz et al., 2013; Keilholz, 2014). More strongly, Bolt et al. (2022) argue that much of the apparent richness of time-varying functional organization reduces to a small number of low-frequency spatiotemporal patterns and traveling waves. Relatedly, Raut et al. (2021) report that spontaneous fMRI fluctuations track global waves that propagate through neocortex, thalamus, striatum and cerebellum in step with arousal, and argue that these waves account for much of the observed functional connectivity structure. If that account is correct, the boundaries between states are placed by the analyst, and the transition matrix is largely a summary of the segmentation procedure.

A third line of work sits between the two pictures. Energy-landscape analyses fit a pairwise maximum-entropy model to fMRI data and describe brain activity as motion in a landscape whose local minima are discrete states, with the passage between basins treated as the object of interest (Watanabe et al., 2014; Ezaki et al., 2018). That description grants discrete states and continuous motion between them at the same time. It does not, however, say how long a passage takes, because the landscape is estimated from how often each pattern occurs rather than from the time course of a crossing.

The distinction between these two scenarios is not semantic. It matters mechanistically: a fast switch implicates competitive, quasi-bifurcation processes with a rapid change in the effective equations of motion, whereas a gradual traversal implicates propagation along a manifold under slowly varying drive. These call for different generative models (Deco and Jirsa, 2012; Khona and Fiete, 2022). It matters for interpretation: if transitions are not events, dwell times and transition probabilities inherit an arbitrary discretization, and their reported relationships to behavior and to clinical status are problematic (Lurie et al., 2020). And it matters for intervention: a growing literature identifies the regions whose stimulation would move the brain from one state to another, or computes the control energy required to do so (Gu et al., 2015, 2017; Deco et al., 2019; Cornblath et al., 2020). That enterprise presupposes that transitions are identifiable events with identifiable drivers.

The question is also genuinely difficult, and has proved contentious in adjacent domains. Whether parietal neurons ramp continuously toward a decision bound or step discretely between states was hampered by inspection of trial-averaged firing rates and benefited from explicit model comparison at the single-trial level (Latimer et al., 2015). The debate that followed (Shadlen et al., 2016; Latimer et al., 2016), and its eventual reformulation within switching state-space models that contain both possibilities as special cases (Zoltowski et al., 2020), is instructive. Averaging destroys the distinction in both directions: a population of sharp transitions with jittered onsets averages to a smooth ramp, and a genuinely gradual process can be made to look abrupt if the alignment marker is placed at the steepest point of change. Any test therefore has to be designed so that both answers remain available.

In resting-state fMRI, “transition” is almost always operationalized as a change in a state label. Hidden Markov Models supply transition probabilities but no description of the dynamics during the switch. Sliding-window methods blur across the switch by construction (Leonardi and Van De Ville, 2015). What has been missing is a framework in which the dynamics on either side of the boundary, and the change in those dynamics at the boundary, are themselves estimated quantities.

Here we use a Switching Linear Dynamical System (SLDS) (Ghahramani and Hinton, 2000; Barber, 2006; Smith et al., 2010; Linderman et al., 2017, 2019; Taghia et al., 2018) for this purpose (Fig. 1A). An SLDS estimates a set of latent variables that capture system dynamics in a space of much lower dimension than the data, a separate linear dynamical system for each state that specifies how the system evolves while in that state, and a probabilistic model of switching between states. Two features make it suited to the present question. First, the model supplies a segmentation that can be treated as an oracle: once time points are labeled, the *measured* fMRI signals around a labeled boundary can be examined directly, and they are free to be smooth or abrupt (Fig. 1B). Second, the model supplies an estimated flow — a vector field — whose change around the boundary can be quantified, and quantified in a graded manner, so that sharpness becomes a measured property of each transition pair.

**Figure 1:**
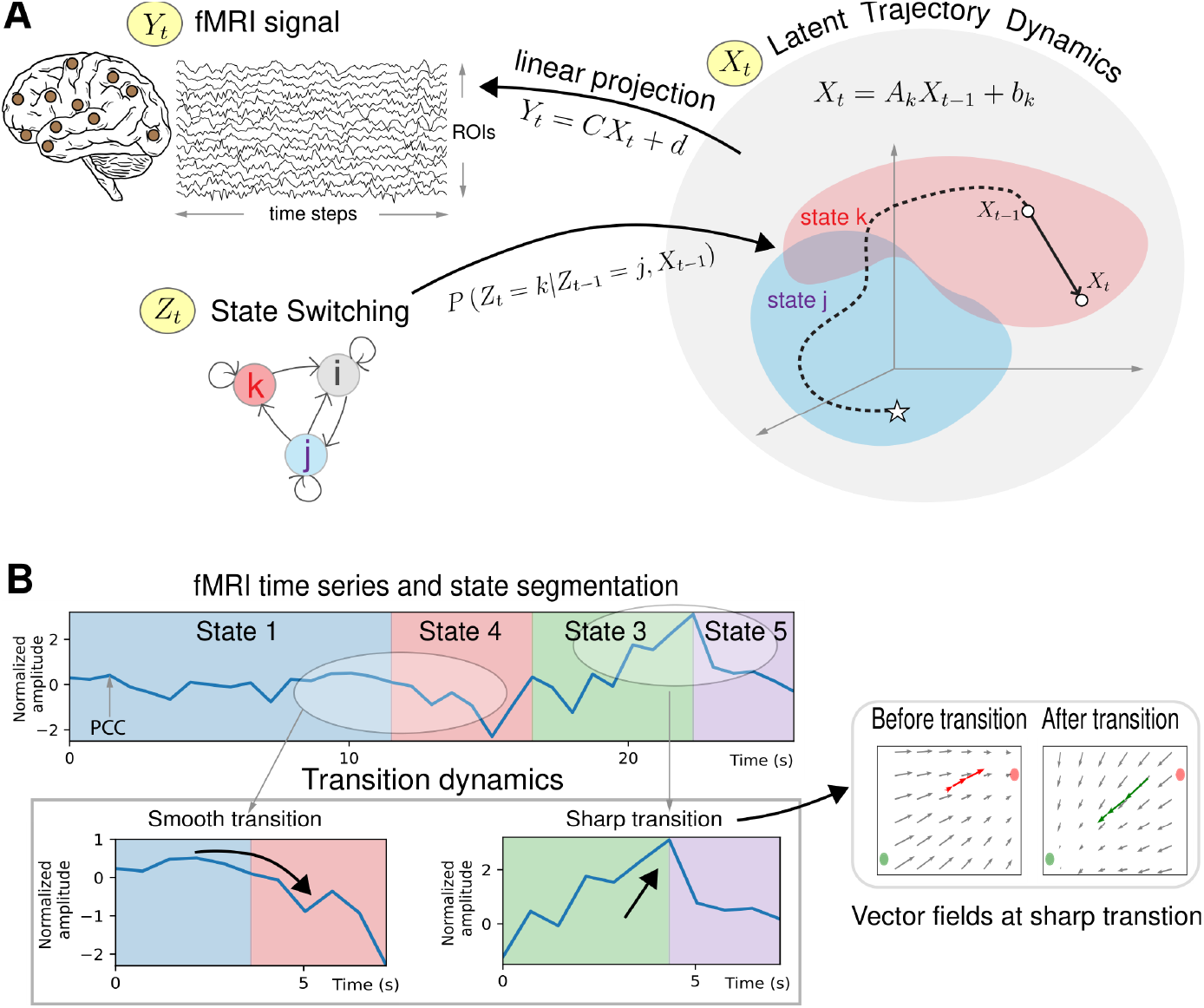
SLDS framework. **A**. A Switching Linear Dynamical System (SLDS) was used to model fMRI resting-state time series data (*Y*_*t*_) from a set of cortical and subcortical brain regions of interest (ROIs). The model represents brain signals in terms of a set of latent variables (*X*_*t*_). For each state *k*, the temporal evolution of the system is specified via a linear dynamical system. In the diagram, the system starts in state *j* (white star) and transitions to state *k*. In state *i*, the system evolves according to the dynamics matrix *A*_*k*_. The model switches between states according to a Markovian process that determines the probability that the system is in state *k* at time *t*. Although the system’s trajectory lives in the latent space, it can be mapped back to the brain space via a linear projection learned from the data. **B**. The SLDS model segments the data into states, allowing us to study the dynamics of the transitions (actual data from the posterior cingulate cortex, PCC). Whereas one transition is visually fairly smooth, the other is considerably sharper, which can be quantified by the change in the vector fields around the time of transition.

The model used here is also a methodological advance. Group analyses of brain-state models are usually built by concatenating participants and fitting one model, so that the states and their dynamics are common to everyone and participants differ only in when each state occurs. We instead fit a *multi-level* SLDS, in which every participant has their own dynamics and transition parameters, tied to group-level parameters by a hierarchical prior and estimated jointly with them in a single pass, following the approach of Linderman et al. (2019) for hierarchical state-space models. The linear projection from latent space to the brain (Fig. 1) is shared, which keeps all participants in a common latent space and makes latent quantities comparable across them.

In the present study, we asked three questions. (1) How does brain activity evolve around state transitions, and is that evolution homogeneous across transition types? (2) Can transition sharpness be quantified in the dynamics themselves, and is any transition sharp enough to warrant the label “toggle”? (3) Which brain regions drive a given transition, and are they the same regions that steer the dynamics within the states involved? For the third question we developed a new region-level measure of *transition importance*, complementing the measure of within-state region dynamics importance we introduced previously (Misra and Pessoa, 2025). Because our parcellation included 54 subcortical regions alongside 200 cortical ones, subcortical contributions were estimated on the same footing as cortical ones.

## Results

### Dynamic states and their transitions

We fit a multi-level SLDS with six states (*K* = 6) and ten latent dimensions (*D* = 10) to Human Connectome Project resting-state data (*N* = 500). To reduce overfitting and increase generalizability, both values were selected on a separate set of *N* = 100 participants not used in any analysis reported below (Methods).

An SLDS model assigns state labels to each time point in a time series. For example, a short segment of fMRI data might be labeled as follows: …33355555116444…, where each time point is associated with a state (note that this short sequence is not representative of our results in which states typically last longer). SLDS states and state transitions are defined in a latent space of dimensionality (D = 10) considerably lower than that of the original brain data space (254 brain regions) (Fig. 1A). At each time point a single state is “active”. The generative model learned from the data estimates a dynamics matrix for each state that specifies how activity evolves during the state; specifically an equation that estimates signals at *t* + 1 based on signals at *t*. Furthermore, based on the theory of linear systems, qualitative properties of temporal evolution depend on properties of the dynamics matrix. For example, the dynamics might be an attractor that converges to a fixed point. Formally, the type of dynamics is given by the eigenvalues of the dynamics matrix. Switching between states follows a Markov process whose transition probabilities are additionally modulated by the current latent position. Because the trajectory lives in a latent space, it is mapped back to the brain through a linear transformation (i.e., emission model) learned from the data.

All six estimated states were stable attractors at the group level (each *p* < 0.05, corrected), so that in the absence of fluctuations in inputs or endogenous factors the system is predicted to settle into a particular configuration. Fig. 2 displays the mean activity of each state at “equilibrium” time (defined as the mean state duration plus three samples). We note briefly that the states were not in one-to-one correspondence with canonical large-scale networks (e.g., default, salience, etc.); to keep this study focused, we will not pursue this relationship here. Across states the SLDS accounted for 43% (State 6) to 57% (State 5) of the variance in one-step-ahead prediction. Activity maps at equilibrium were positively correlated among State 1, State 2 and State 3 and anticorrelated with State 4, State 5 and State 6 (Fig. S1), with the strongest anticorrelations involving the default-related State 3 and the attention/control-dominant State 5. Transitions were structured rather than random. Of the 30 possible directed transitions, only 10 were statistically significant by a group permutation test (Fig. S2). State 1, which was occupied roughly half the time, transitioned reliably only to State 4 and State 5, while State 2, State 4 and State 5 reliably transitioned to State 1. Several transitions were unidirectional: notably, the attention/control-dominant State 5 transitioned to the default-related State 3, but not the reverse; State 3 instead transitioned to the sensorimotor-dominant State 4. The remainder of this paper concerns what happens during such state transitions.

**Figure 2:**
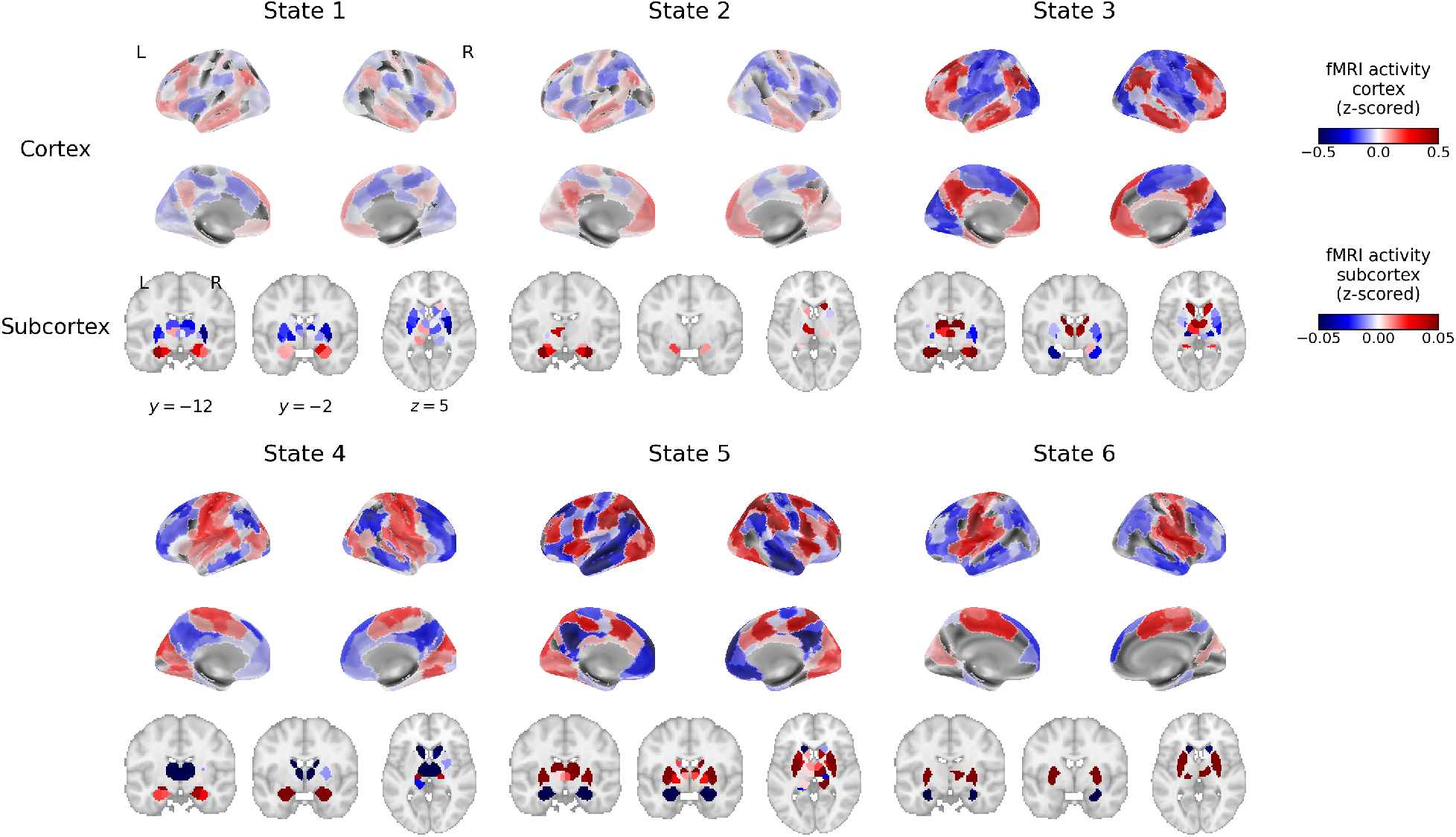
State activity maps. fMRI activity was determined based on mean activity at the region (ROI) level at “equilibrium” time (mean state duration plus three time steps). Activity was normalized (z-scored) for each region separately in each run; thus, positive and negative values should be interpreted relative to a region’s overall temporal mean. States are ordered by their mean duration. Maps are thresholded based on False Discovery Rate at 0.05 (6 states, 254 whole-brain regions). MNI coordinates are shown.

### Signal evolution around state transitions is heterogeneous

We first examined the measured fMRI signal in temporal windows centered on state transitions (Fig. 3; all 10 significant transitions in Fig. S3). This analysis uses the model only to place the boundary; the shape of the activity around that boundary is not constrained by the model and could have come out flat, monotone, or reversing.

**Figure 3:**
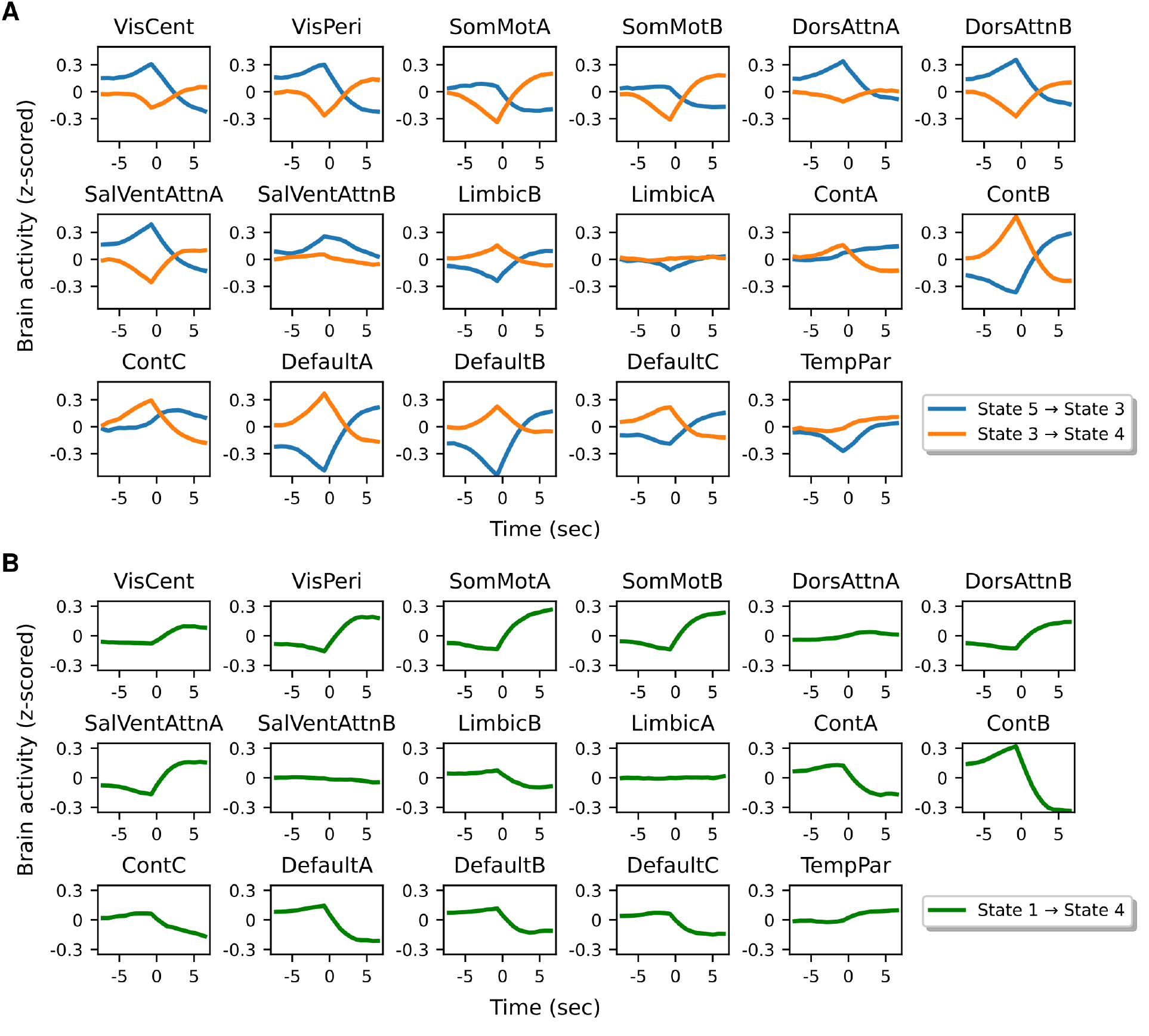
Network activation changes around state transitions. **A**. Two transitions with sharp, toggle-like profiles: several canonical Yeo-17 networks reverse the sign of their activity change across the boundary. **B**. A smooth, “one-sided” transition: activity is close to flat before the boundary and changes only afterwards. *t* = 0 indicates the first time point of the new state. Line thickness indicates standard error of the mean across participants. All significant transitions are shown in Fig. S3.

Two transitions describe one extreme (Fig. 3A). To illustrate activity changes, we mapped the 200 cortical ROIs to Yeo-17 networks for labeling purposes and averaged signals across ROIs belonging to a given network. For State 5 ↦ State 3, DefaultA/B/C activity was decreasing before the transition and increasing after it, while DorsAttnA/B did the opposite; several networks thus reversed the sign of their signal evolution across the boundary. The same held for State 3 ↦ State 4, with SomMotA/B increasing and DefaultA/B/C decreasing. The clearest reversals were in DefaultA and ContB, whose curves for the two transitions cross around *t* = 0.

State 1 ↦ State 4 illustrates the other extreme (Fig. 3B). Here activity was close to flat before the transition and changed only afterwards, in the same direction, for most networks — a “one-sided” profile. Inspection of all significant transitions (Fig. S3) showed that a preponderance of toggle-like reversals characterized State 3 ↦ State 2, State 3 ↦ State 4, and State 5 ↦ State 3, whereas one-sided profiles dominated State 1 ↦ State 4, State 1 ↦ State 5, and State 2 ↦ State 1, with the latter group nevertheless showing sharp reversals in individual networks (for example, ContB during State 1 ↦ State 4).

Two features of this comparison deserve emphasis. First, the transitions are averaged across instances and participants, and any jitter in the inferred boundary blurs the average. Averaging therefore biases the result toward smoothness, making the observed reversals conservative. Second, sharp and smooth transitions were not segregated by the networks involved. State 1 ↦ State 4 and State 3 ↦State 4 share a destination state and both involve prominent somatomotor engagement, yet differ markedly in profile; State 2 ↦ State 1, State 4 ↦ State 1 and State 5 ↦ State 1 also share a destination state and also differ. Regional differences in hemodynamic response shape are therefore unlikely to account for the heterogeneity.

### Quantifying change in dynamics at a state transition

The central goal of our study was to characterize state transitions. In particular, we sought to test whether some transitions were more toggle-like, signaling a *change of dynamical regime*, and others smoother, signaling a reorientation within a similar regime. The analyses above describe the fMRI *signal* around a transition but not how abruptly the *dynamics* change at the boundary.

We therefore turned to the vector field defined by the system (Fig. 1B): at every position in latent space, a vector giving the direction and magnitude of the expected one-step evolution (a particle placed at that position moves along that vector in one time step). The field was estimated empirically, from one-step differences of the inferred latent trajectory within local neighborhoods, separately at each time point relative to the transition (Methods). Two properties of this estimate matter for what follows. It is computed from the trajectory the system actually followed, and it is estimated separately at each time step so it is free to change gradually.

Fig. 4 shows fields projected onto two latent dimensions for two transitions (all significant transitions in Fig. S6). Consider State 3 ↦ State 4. For several time steps before the boundary the field pointed toward the attractor of the current state (red) and the system moved in that direction. At *t* = 0, the crossing step, the field *flipped*, and thereafter it pointed toward the attractor of the new state (green). State 5 ↦ State 3 behaves the same way: the field before the boundary carries the system toward the attractor of State 5, and from the crossing step onward toward the attractor of State 3.

**Figure 4:**
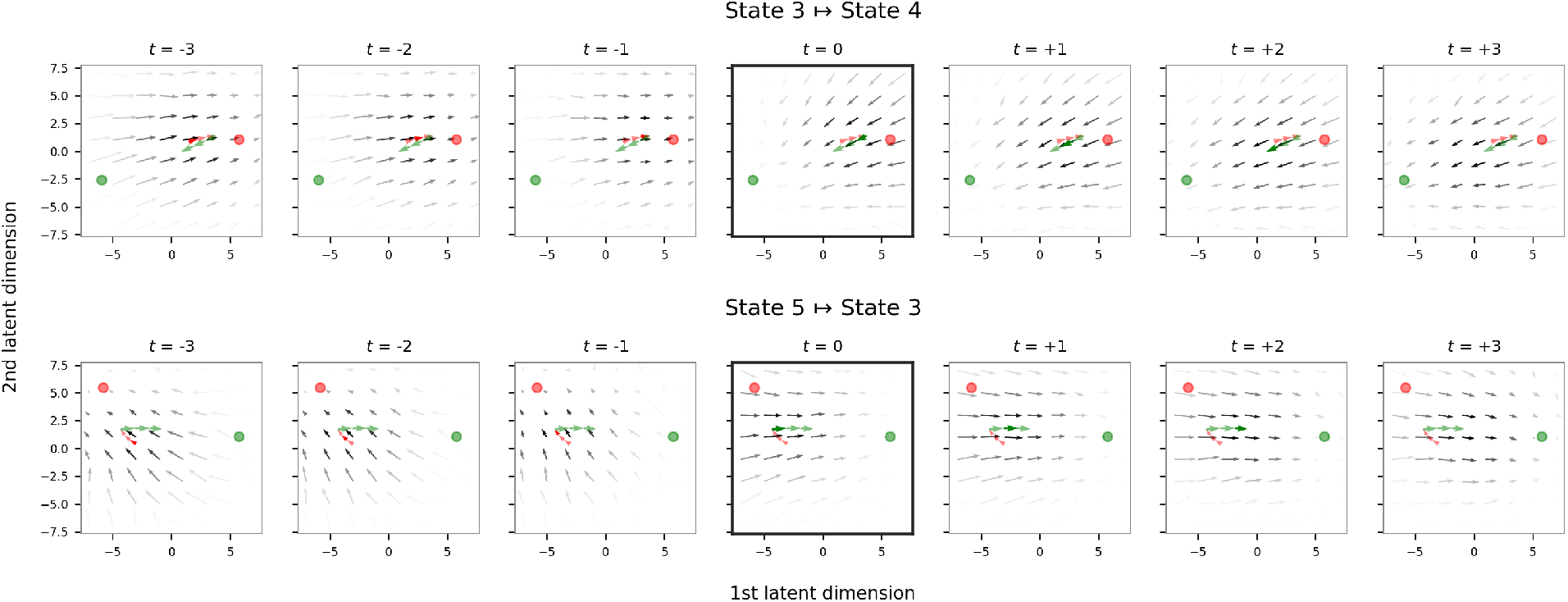
Quantifying the change in dynamics at a state transition. Vector fields estimated from one-step changes in latent state and projected onto two dimensions for visualization. Each panel is one time step; a displacement is labeled by the frame it arrives at, so *t* = 0 is the step that crosses the boundary, from the last sample of the old state to the first sample of the new one (Methods). Red/green vectors represent the mean trajectory during the transition window (red: original state; green: new state). Red and green dots represent the point attractors. The grayscale values of the vectors are proportional to the number of data points that fall within the associated neighborhood, indicating data density at time *t*. Fig. S6 displays all statistically significant transitions.

### Does the flow reverse?

To quantify state transitions, we created a *transition sharpness* index defined as the cosine similarity between the fields at adjacent time points (Fig. 1B). A negative value means that the estimated flow reversed direction between two samples acquired 720 ms apart. This is the natural instrument for the question we are asking for three reasons. It is a statement about direction alone and is unaffected by how fast the system happens to be moving, so it cannot be driven by one state being slower than another. It is resolved in time, so it says not only that the flow changed but when. And it inherits the two properties of the field estimate described (i.e., described from the trajectory the system actually followed; estimated at each time step), so it is *not* imposed by the model’s discrete architecture.

Transition sharpness was determined in the full ten-dimensional latent space (Methods; Fig. 5). Because a cosine between two noisy averages is pulled toward zero, each transition was also given a floor: the identical estimator applied to pseudo-transitions taken from the middle of long dwells of the same two states, so no change of rule occurs at a pseudo-transition. Two of the ten significant transitions, State 6 ↦ State 2 and State 6 ↦ State 5, are set aside here and in everything that follows. The floor requires eleven consecutive frames carrying a constant state label, and State 6, whose mean dwell is 2.9 s, cannot supply them. The measure is therefore reported for the remaining eight transitions. The results (Fig. 5) followed the same shape in every case. Away from the boundary the cosine sits between 0.64 and 0.96, close to the transition’s own floor, and it drops to a single-sample trough at *t* = 0. Four transitions reversed: State 3 ↦ State 4, State 5 ↦ State 3, State 4 ↦ State 1 and State 5 ↦ State 1. State 3 ↦ State 2 fell to +0.09, while State 1 ↦ State 4, State 2 ↦ State 1 and State 1↦ State 5 remained clearly positive, a reorientation without reversal.

**Figure 5:**
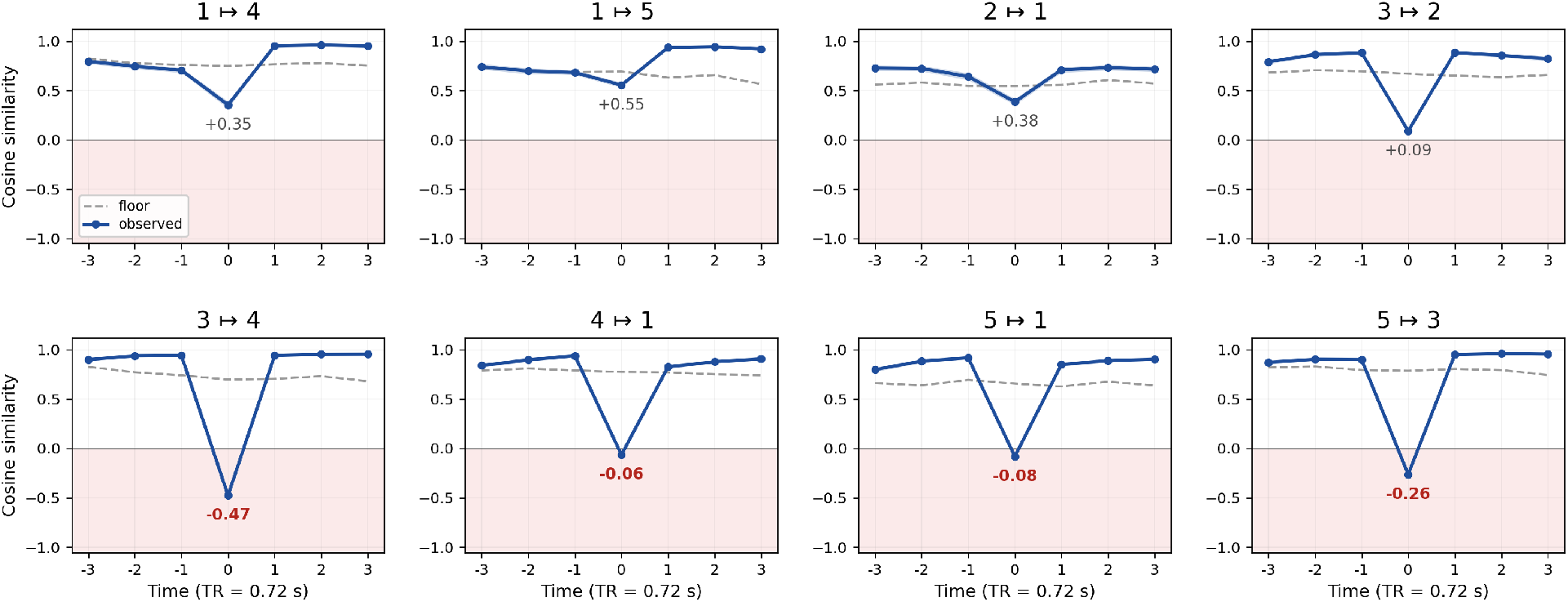
Does the dynamics flow reverse? For each of the transitions, the cosine similarity between the flow field at time *t* and the field one time step earlier, with neighborhoods defined by *k*-means in all ten latent dimensions (blue; shaded band is the 95% confidence interval from subject subsampling). As in Fig. 4, a displacement is labeled by the frame it arrives at, so *t* = 0 is the step that crosses the boundary. Values in the shaded region below zero indicate that the estimated flow reversed direction between two samples acquired 720 ms apart; the value at the crossing step is printed in each panel, in red where the confidence interval lies entirely below zero. The dashed gray curve is the transition’s own reference floor: the identical estimator applied to pseudo-transitions in the middle of long dwells of the same two states, with the number of contributing points matched cell by cell and lag by lag. Bonferroni correction across 8 tests.

Note that the reversals are not symmetric between the two directions of a pair. Neither exit from the high-occupancy State 1 reversed the flow, at +0.35 for State 1 ↦ State 4 and +0.55 for State 1 ↦ State 5, whereas two of the three entries into it did, at −0.06 for State 4 ↦ State 1 and −0.08 for State 5 ↦ State 1; the third entry, State 2 ↦ State 1, did not, at +0.38. Whatever distinguishes a sharp transition from a smooth one is therefore not a property of the pair of states alone, but of the direction in which the pair is traversed.

The analyses above provide strong support for the existence of toggle-like transitions in the human brain as detected with functional MRI. To test the robustness and generality of our finding, we proceeded as follows. Transition sharpness was determined in the latent space. We reasoned that if the latent representation is a valid description of brain activation, we should obtain evidence of the transitions and their sharpness based on the original fMRI signal space, *Y*. Indeed, as described in Appendix B, the transition sharpness based on fMRI followed almost exactly the same ranking as the one in latent space. Evidence for toggle-like transitions in Fig. 5 was obtained with a model with six states. In supplementary analyses (Appendix C), we fit the data using *K* = 8 states and evaluated transition sharpness. As before, we observed transitions of varying smoothness, including multiple that were toggle-like, where the field reversed at state transitions (but note that with 4 states no reversals were detected, although varying smoothness was observed). We also detected toggle-like transitions in supplementary analyses with data that did not remove the global signal (Appendix C). Taken together, these analyses further strengthen the evidence for the existence of sharp transitions in human fMRI resting data.

### How much do the vector fields differ?

The cosine measure of transition sharpness determines whether the flow turned round. It does not say how much of the system’s motion was altered in turning. We addressed that second question with the *field contrast*: the distance between the vector fields of the origin and destination states, evaluated at the points the system actually traverses around the transition and averaged over them (Methods). It is computed in the full ten-dimensional latent space.

Across transitions, the field contrast ranged from 1.24 for State 2 ↦ State 1 to 2.11 for State 5 ↦ State 3, the largest exceeding the smallest by about 70%. Its two largest values, State 5 State 3 and State 3 ↦ State 4, belong to the two transitions whose flow reverses most deeply. The two measures therefore agree about which transitions alter the dynamics most.

Away from those extremes they agree only moderately. The rank correlation between the field contrast and the cosine index was *ρ* = −0.57 (*n* = 8, exact *p* = 0.15). The reason follows from what the field contrast measures. A distance between two vectors is symmetric for the two states it compares; the field contrast is blind by construction to the entry-versus-exit asymmetry reported above. Thus, the two measures are best read as answering different questions rather than as corroborating one another, and we rely on the cosine for the claim that some transitions are toggle-like.

One caveat belongs here rather than in the Discussion. Both measures contrast two states’ fields and are nonzero by construction whenever the states differ, so neither absolute value carries meaning on its own. What is interpretable is the spread across transition pairs. That spread is recovered by two analyses with different dependence on the model: the measured network time courses (Fig. 3), which use the model only for segmentation, and the time-resolved fields (Fig. 4), which use the inferred trajectory but not the state dynamics matrices. The field contrast adds a third view, and it agrees with the other two about the extremes.

### Which regions drive a transition?

How strongly is the brain driven toward a switch, and when? At each moment, the model establishes how far the current whole-brain pattern points along the direction that favors state *j* over staying in state *i*. We call this the *drive* toward that switch. We evaluated it on the fMRI activity at each frame separately, so that it uses no information from any other time point (Methods). In the middle of a dwell (the midpoint of a state) it is essentially zero (leaving the odds of a switch at their usual value, the 1 line in Fig. 6). Over the four seconds before a state boundary it rises steadily, reaching its maximum on the last frame of the departing state. At that maximum the odds of that specific transition are between 1.8 and 3.2 times their mid-dwell value, largest for State 3 ↦State 4 at 3.2 and State 5 ↦ State 3 at 2.5. It then falls back over the following seconds. Two qualifications are in order. The drive being elevated near a boundary is not by itself independent evidence that transitions are events, because the model places boundaries using this very quantity; what is not built in, and what the data determine, is that the drive sits at zero in mid-dwell, so that the brain is pushed toward no particular successor; that the buildup takes several seconds rather than appearing abruptly; and that it is discharged rather than sustained afterwards. And the size of the effect (e.g., a doubling to tripling of the odds) is not property of the fit.

**Figure 6:**
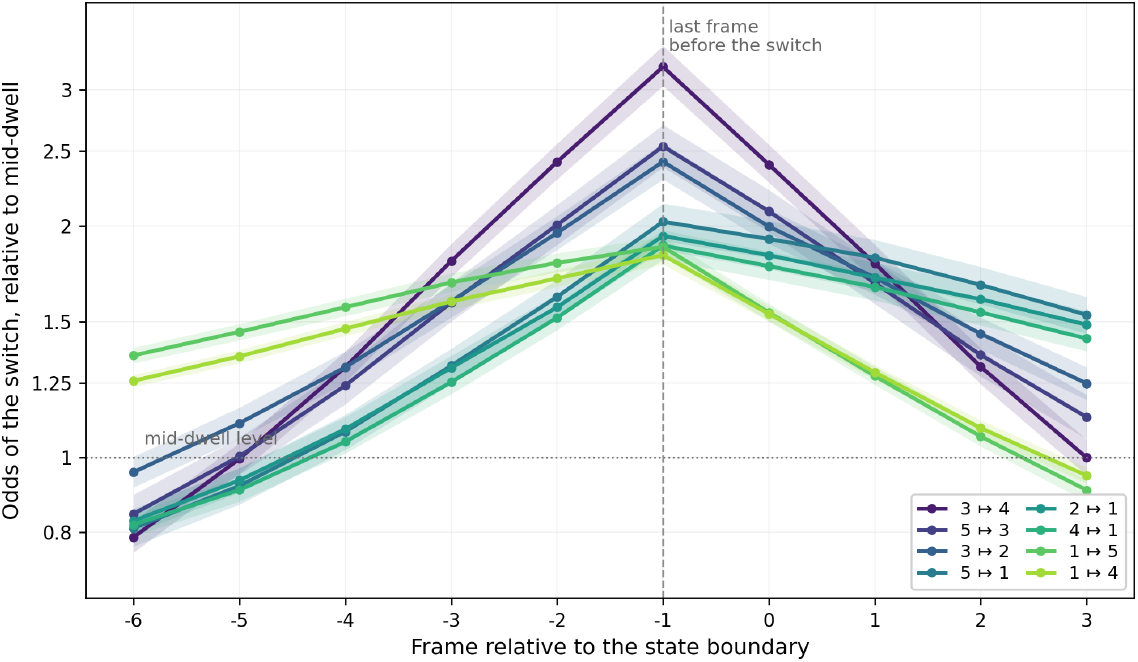
The drive toward a state switch. For each of the transitions, the drive (over all regions) expressed as the odds of that switch relative to its mid-dwell value. Frame −1 is the last time point of the departing state. Each curve is referenced to its own mid-dwell level, measured at the midpoints of dwells; shaded bands are group level 95% bootstrap confidence intervals over the 500 participants. The axis is logarithmic in the odds, which makes the region shares of Fig. 7 additive.

The dynamics modeled by SLDS live in a latent space, and the states themselves are inherently multi-variate. We hypothesized that some brain regions contribute to a state transition more than others. To recover the contribution of individual regions we developed a lesion-type analysis targeted at the SLDS transition model (Methods). The logic is as follows. The SLDS model estimates the odds of leaving state *i* for state *j* from the brain’s current configuration, which every region helps to specify. Removing the contribution of a single region *m* yields a counterfactual estimate of where the system is *without letting region m inform it*. The resulting change in the odds of the *i*↦ *j* switch defines the *transition importance* of region *m* for that switch. Because signals contributing to a transition must precede it, importance maps were computed over the three samples preceding the state boundary.

Because the drive is a sum over regions, each region’s contribution is a share of it. The highest-ranked region accounted for between 1.4% and 3.6% of the drive toward its transition, which is relatively modest. Ten regions, however, carried between 13% and 24% of it depending on the transition, 22% for State 3 ↦ State 4 and 21% for State 5 ↦ State 3 (Fig. 7). About 5% of the parcellation therefore supplied around a fifth of everything pushing the brain toward a given switch.

**Figure 7:**
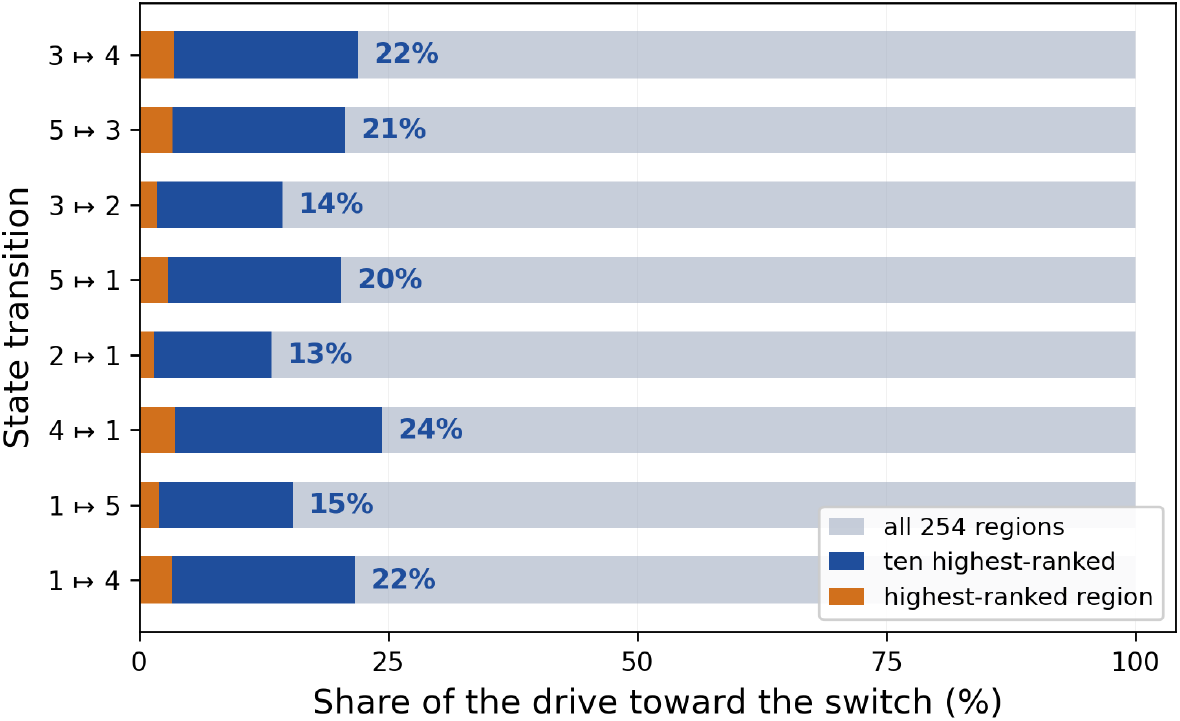
How much of the drive individual regions account for. For each of the transitions, the share of the drive in the three time points before the boundary contributed by the single highest-ranked region and by the ten highest-ranked regions, out of all 254. Shares are computed from the same instances and the same window as Fig. 6. Transitions are ordered as in Fig. 6.

Fig. 8A shows importance maps for two transitions (all significant transitions in Fig. S4). For State 5 ↦ State 3 the strongest contributions came from default-mode regions, in particular the angular gyrus and posterior cingulate cortex. For State 3 ↦ State 4 the most important regions belonged to networks ContA and ContB, with a notable contribution from a DefaultA region as well. Thus different transitions out of, or into, structurally overlapping state configurations recruited distinguishable sets of regions. Notably, in the State 5 ↦ State 3 transition the most important regions were default-mode regions, and State 3 itself exhibited strong overlap with the default-mode networks, while State 5 overlapped with task-positive networks. For this transition, then, the regions carrying most influence over the switch were those characteristic of the state being entered (i.e., the future state) rather than of the state being left.

**Figure 8:**
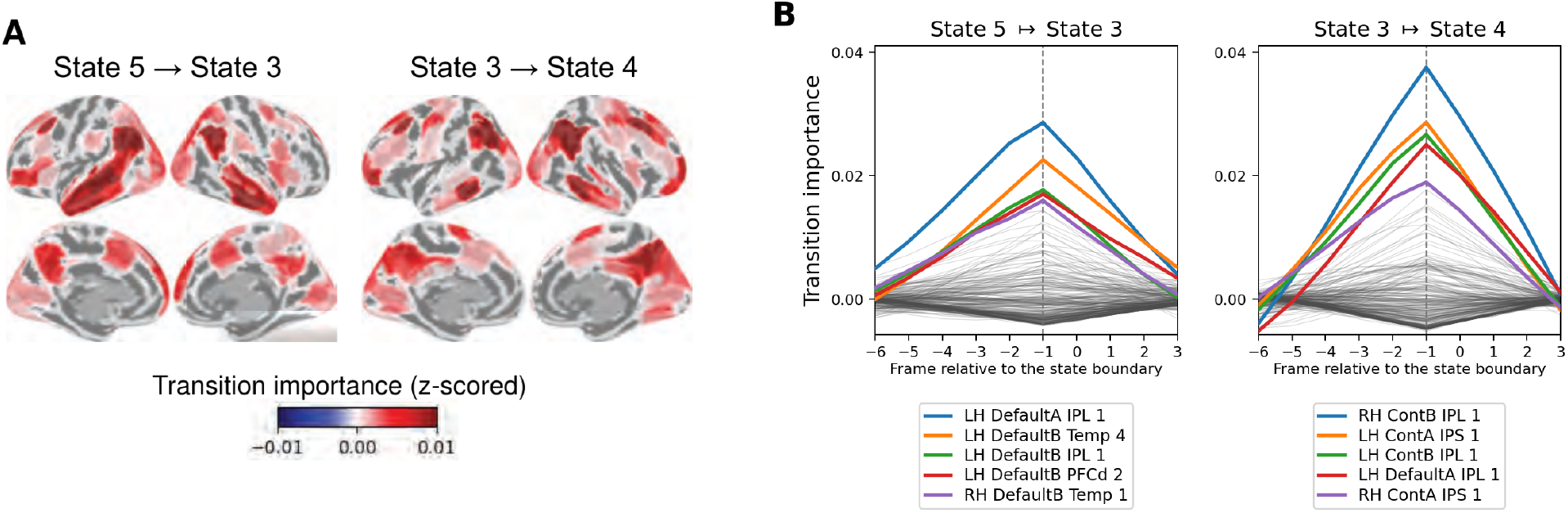
State transition importance. **A**. Importance maps for two transitions with sharp transition profiles. Maps thresholded against the null of equal importance across regions (False Discovery Rate at 0.05). The two transitions recruit distinguishable sets of regions: default-mode regions for State 5 ↦ State 3 and control-network regions for State 3 ↦ State 4. **B**. Importance time series around the transition, on the same frame axis as Fig. 6: frame 0 is the first sample of the new state and the dashed line marks frame −1, the last sample of the departing state. Each curve represents a brain region, with the top five highlighted. Importance rises steadily before the boundary, peaks on the last frame of the departing state, and falls immediately thereafter. The complete set of transition importance maps and importance time series for significant transitions are shown in Figs. S4 and S5.

The temporal profile of transition importance was equally informative (Fig. 8B; all transitions in Fig. S5). Computed in a sliding window, importance for the top-ranked regions rose steadily over the seconds preceding the boundary, peaked on the last frame of the departing state, and fell immediately thereafter. This profile was observed for all eight transitions investigated, and it is the regional counterpart of Fig. 6: the buildup and discharge established there for the whole brain was observed at the level of individual regions.

### Regions that steer a state and regions that end it

Transition importance was developed here to quantify a region’s relative contribution to state change. In this section, we compared it directly against the measure of *within-state* region dynamics importance we developed previously (Misra and Pessoa, 2025), which quantifies how much the predicted one-step evolution of whole-brain activity changes when a region’s contribution is removed from the state’s dynamics matrix. The two measures ask different questions of the same model: which regions steer the trajectory while the system is in a state, and which regions push the system out of it.

Here, we ask the question of the relationship of these two types of region importance. Consider State 5 (Table S1). Within-state dynamics were dominated by dorsal attention and visual regions, consistent with the attention/control character of that state (see also Fig. 2). The regions that drove the State 5 ↦ State 3 transition, however, were default-mode regions, which were not prominent in State 5’s within-state dynamics. In other words, default-related regions accumulated influence over the course of an attention/control state and eventually terminated it, without being the regions that governed its internal evolution. Such dissociation is not universal. For State 3 ↦ State 4, control-network regions in the intraparietal sulcus and inferior parietal lobule contributed to the transition and were also among the most important regions for State 3’s within-state dynamics. Table 1 shows results for all eight transitions considered showing that the amount of overlap varies across transitions, with some of them exhibiting a pattern of dissociation between within and between region importance. For the two exits from State 1, the ten regions that steer the state have average ranks of 236 and 209 (of 254) for ending it, well past the 127.5 expected by chance, so the regions that hold the baseline regime together are among the last to break it (i.e., help transition from it). For State 3 the two sets largely coincide, at average ranks near 34 with three or four regions shared outright, so whether steering and ending are separable depends on the regime rather than on the measure.

**Table 1:** Do the regions that steer a state also end it? For each transition, the 254 regions were ranked twice. The first ranking uses within-state region dynamics importance (Misra and Pessoa, 2025), which measures how much the predicted one-step evolution of whole-brain activity changes when a region’s contribution is removed; its ten highest-ranked entries are the regions that most *steer* that state. The second ranking uses transition importance for that specific switch (with rank 1 the region contributing most to the switch); its ten highest-ranked entries are the regions that most help *end* the state. The second column counts the regions common to the two top tens. The third column takes the ten steering regions and gives their mean rank in the transition importance ranking, so that a small value means the regions steering the state are also prominent in ending it, and a large value means they are among the last regions to end it. For example, the ten regions that steer State 5, listed in Table S1, occupy ranks 24 to 77 of 254 for ending it, an average of 46. Both rankings are averages over the 500 participants. Ten regions drawn at random average rank 127.5, and two unrelated top-ten lists share 0.39 regions on average and share none 66% of the time, so a zero in the second column is not by itself informative and the third column helps separate the cases. The relationship follows from the third column against a permutation null over regions: *opposed* when the steering ten fall near the bottom of the ending ranking, *unrelated* when they sit near the middle, and *shared* when they sit near the top. Transitions involving the rare state, State 6, are excluded, as elsewhere.

| Transition | Shared of top ten | Average rank of the steering ten | Relationship |
| --- | --- | --- | --- |
| STATE 1 $\mapsto$ STATE 4 | 0 | 236 | opposed |
| STATE 1 $\mapsto$ STATE 5 | 0 | 209 | opposed |
| STATE 5 $\mapsto$ STATE 1 | 0 | 121 | unrelated |
| STATE 4 $\mapsto$ STATE 1 | 0 | 82 | shared, weakly |
| STATE 5 $\mapsto$ STATE 3 | 0 | 46 | shared |
| STATE 2 $\mapsto$ STATE 1 | 2 | 32 | shared |
| STATE 3 $\mapsto$ STATE 4 | 4 | 34 | shared |
| STATE 3 $\mapsto$ STATE 2 | 3 | 34 | shared |

## Discussion

Studies of the resting state collectively indicate that it should be understood dynamically. Here we asked how the brain moves between resting-state configurations, and in particular whether that movement is toggle-like or gradual. The answer is that it is both, depending on which pair of states is considered. Three analyses support this conclusion, and they differ in how much they depend on the SLDS model. The measured activity of canonical networks around a state boundary reversed sign for some transitions and was flat before the boundary for others. The flow field estimated in latent space reversed direction between two samples acquired 720 ms apart for some transitions and merely reoriented for others. And the field contrast, a full-dimensional measure of how different the equations of motion of the origin and destination states are, varied across transitions by about 70 \% from smallest to largest,and agreed with the other two about which transitions alter the dynamics most. A new lesion-type measure then identified the regions that drive individual transitions, and showed that their influence accumulates over the seconds before the switch and dissipates immediately after it.

### State transition sharpness: from toggle-like to smooth

We sought to test the hypothesis that, if brain states are a useful way to conceptualize fMRI activity during the resting state, transitions between them should be “non trivial”. Four of the eight transitions reversed the estimated flow between two adjacent samples acquired 720 ms apart: State 3 ↦ State 4, State 5 ↦ State 3, State 5 ↦State 1 and State 4 ↦State 1. Each cleared Bonferroni correction over the eight tests, the weakest at |*z*| = 3.9. Two of the four were particularly pronounced: for State 3 ↦State 4 and State 5 ↦State 3 the cosine fell to −0.47 and −0.26, canonical networks reversed the sign of their activity change across the boundary, and the field contrast was largest. We called transitions of this type *toggle-like*. Multiple toggle-like transitions were also observed in supplementary analyses employing eight states, showing that this behavior did not depend on the particular six-state partition we utilized (Appendix C). Therefore, we propose that a description of fMRI resting-state dynamics in which all apparent state boundaries result from the modeling approach is incomplete. Our results thus help qualify a more continuous picture. Quasi-periodic patterns and low-frequency traveling waves account for a substantial share of time-varying functional organization (Majeed et al., 2011; Keilholz et al., 2013; Bolt et al., 2022), and our own smooth transitions (State 1 ↦State 4 and State 1 ↦ State 5, the two exits from the high-occupancy State 1) are entirely consistent with such a description. But our results suggest it is not the entire story.

At the same time, BOLD is a low-pass filtered, slowly evolving signal, and the fastest change we can resolve is set by the sampling rate (TR = 0.72 s) and by the hemodynamic response, not by the underlying dynamics. Naturally, we cannot determine how fast the neural switch is; we can only say that at the fMRI timescale some transitions are as sharp as the measurement permits while others are not. Because sharp and smooth transitions shared destination states and engaged overlapping networks (most directly, State 1 ↦ State 4 and State 3 ↦ State 4 both terminate in a somatomotor-dominant state), the heterogeneity cannot be assigned to regional differences in hemodynamic properties (Handwerker et al., 2004). For additional control analyses ruling out hemodynamic explanations, see Appendix D.

What is the importance of transition sharpness? Mechanistically, a reversal of the flow between adjacent samples is what one expects from a competitive process in which the effective equations of motion change quickly: mutual inhibition between configurations, gating, or passage near a bifurcation. It is not what one expects from drift along a manifold under slowly varying dynamics. The two classes call for different generative models, and the empirical demonstration that both occur within a single resting-state session constrains that modeling (Deco and Jirsa, 2012; Cabral et al., 2014; Khona and Fiete, 2022). Practically, sharpness justifies the standard vocabulary of states. Dwell times and transition probabilities are interpretable as properties of the brain to the extent that the boundaries they are computed from correspond to events; where they do not, they are properties of the segmentation (Lurie et al., 2020). Our results suggest that this behavior is transition-specific and should not be assumed globally. Finally, the interventional program that asks which regions to stimulate in order to move the brain from one state to another (Gu et al., 2015, 2017; Deco et al., 2019; Cornblath et al., 2020) is best posed for those transitions that are more pronounced.

### What distinguishes a sharp transition from a smooth one?

Having established that transitions differ, the natural question is what the difference reflects. It is not based on the distance between the states being joined. If it were, the transitions joining the most dissimilar activity configurations would be the sharpest but they are not: State 3 ↦ State 4 joins the most strongly anticorrelated pair (*r* = −0.91) and reverses the flow most deeply, but State 1 ↦ State 5 joins a pair almost as strongly anticorrelated (*r* = −0.80) and is the smoothest transition measured; State 3 ↦ State 2, which joins two *positively* correlated configurations (*r* = 0.39), falls between them.

We suggest that the *direction* of travel with respect to State 1 is an important factor. State 1 was occupied roughly half the time and resembles the long-dwell, high-occupancy “baseline” or hub-like regimes reported elsewhere (Song et al., 2023; Chen et al., 2022); it transitioned reliably only to State 4 and State 5. Leaving the state did no reverse the flow: State 1 ↦ State 5 at +0.55 was the highest cosine measured and State 1 ↦ State 4 at +0.35 was among the highest; these are both cases of a reorientation without reversal. Entering the state usually did: two of the three entries into State 1, State 5 ↦ State 1 and State 4 ↦ State 1, reversed the flow.

We thus suggest that the brain *drifts out* of the baseline regime and is *switched into* it, and that the toggle-like character of resting-state dynamics is concentrated in entries to the baseline regime and in transitions among the more differentiated regimes, such as default-related, attention/control-related, sensorimotor. The potential emerging picture is as follows: escape from a broad, strongly occupied attractor is slow (and possibly noise-driven), whereas convergence into it can be fast. This conjecture predicts that in a model with a finer state repertoire exits from the highest-occupancy state should not reverse the flow, while entries into it may. The eight-state model analysis (Appendix C) is consistent with this idea. There, neither exit from the dominant state there reversed the field, while of the four entries one does.

We also note that the two sharpest transitions both involve the default-related State 3, once as destination (State 5 ↦ State 3) and once as origin (State 3 ↦ State 4). Entries into and exits from the default-related state were thus the most abrupt events we observed. The antagonism between default-mode and attention/control systems is among the most reproducible findings in resting-state fMRI (Fox et al., 2005), and a competitive relationship of that sort would be anticipated to generate fast switches rather than gradual evolution. We are cautious here because global signal regression was applied during preprocessing and is known to affect anticorrelation structure (Saad et al., 2012; Chai et al., 2012). We note that the model fit without global signal regression still produced transitions that reverse the flow (Appendix C), so the phenomenon does not depend on that step.

### Relation to prior work on time-varying functional organization

Several approaches support the view that resting fMRI signals alternate among a small number of whole-brain patterns. Sliding-window clustering revealed recurring connectivity states with non-random transitions and behaviorally relevant dwell times (Allen et al., 2012; Damaraju et al., 2014). Hidden Markov modeling showed that large-scale activity switches among a handful of metastates organized hierarchically in time (Vidaurre et al., 2017, 2018a; Song et al., 2023). Frame-wise co-activation analyses decompose BOLD time series into transient maps with characteristic temporal structure (Liu et al., 2018; Peng et al., 2023; Karahanoğlu and Van De Ville, 2015; Gutierrez-Barragan et al., 2024; Lee et al., 2024). Quasi-periodic pattern analysis identifies repeating spatiotemporal waveforms that explain a substantial share of resting functional connectivity structure (Keilholz et al., 2013; Keilholz, 2014), and recent work shows that much of that structure reduces to a small number of low-frequency spatiotemporal patterns and traveling waves (Bolt et al., 2022; Raut et al., 2021). These approaches have been applied to the same data and compared directly, and they do not produce identical state repertoires (Maltbie et al., 2022).

Our state repertoire is consistent with this body of work, and our finding that transitions are non-random reproduces one of its central results. Importantly, the present analysis adds to this literature a quantitative characterization of *boundaries*. Each of the methods above yields state labels, but do not estimate the dynamics on either side of a boundary. So they cannot determine the type of change experience at the boundary. That is precisely what the SLDS modeling framework provides, and allowed us to test the toggle-versus-smooth question. The finding that both regimes exist suggests that the discrete and continuous descriptions in the literature are not competitors so much as descriptions of different parts of the broader dynamic repertoire during the resting state.

A word about the model developed here. Every quantity reported above is an average over 500 participant-level fits rather than a summary of one concatenated time series. This follows from the multi-level formulation: subject-level dynamics are estimated with partial pooling toward the group, which is what makes per-participant dynamics estimable at all from roughly 4,800 frames each, while the shared emission model (i.e., latent state to brain linear transformation) keeps the latent space common so that vector fields and their angles can be compared across people. The cost of that choice is the assumption it rests on, namely that the mapping from latent space to regional activity does not vary across participants; relaxing it, and allowing individual differences in the emission model as well as in the dynamics, is a natural extension for future work.

### Region contributions to transitions

The multivariate SLDS approach leaves open the question of which brain regions impact state transitions. The transition importance measure introduced here addresses this question explicitly. The definition employed is counterfactual: instead of asking how much a region’s activity changes, it asks how much the model’s evidence for a particular switch depends on considering a region’s signal.

We highlight two results here. First, transitions had identifiable and distinguishable regional drivers. Default-mode regions, chiefly angular gyrus and posterior cingulate cortex, drove State 5 ↦ State 3, whereas control-network regions in intraparietal sulcus and inferior parietal lobule drove State 3 ↦ State 4. Second, importance had a stereotyped time course: a steady rise before the boundary, a peak at the boundary, and an immediate fall. It is interesting to note that this is a “precursor signature” resembling early-warning indicators that precede regime shifts in other complex systems (Scheffer et al., 2009). Of course, determining critical slowing down or any specific bifurcation is particularly challenging given the hemodynamic timescale.

In our view, the dissociation between within-state dynamics importance and transition importance is the most relevant region-level result. In five of the eight transitions, the ten regions that steer the origin state’s internal evolution and the ten that drive the switch out of it share no region at all. How separable the two roles are depends more on the state being left than on the state being entered, though not without exception: the two exits from State 1 behave alike and the two from State 3 are nearly identical.

State 1 and State 3 mark the extremes, and they differ in anatomy as well as in degree. The internal dynamics of the high-occupancy State 1 are steered almost entirely from the striatum and thalamus, nine of its ten highest-ranked regions being subcortical, while the regions that end those transitions are cortical. In State 3 the same control regions in the intraparietal sulcus and inferior parietal lobule serve both roles, with four of the top ten common to the two lists. Exits from State 5 and State 4 fall in between, sharing no region while the steering set still sits in the upper half of the ending ranks.

We draw two conclusion here. Maintaining a regime and ending it are separable functional processes, and how separable they are is itself a property of the regime. More generally, we suggest that activity magnitude, the quantity almost universally mapped in fMRI, does not uniquely determine a region’s contribution to system-level dynamics. Functional MRI focuses on amplitude, and a region that changes its activity is interpreted as being involved. Neither of the two measures compared here is a measure of amplitude. Within-state dynamics importance asks how much the predicted evolution of whole-brain activity changes when a region is removed from the state’s dynamics matrix, and transition importance asks how much the model’s evidence for a particular switch depends on that region’s signal. In both cases a region enters through the alignment of its activity with a direction in the latent space, so a region can be strongly modulated and contribute little, or contribute substantially while being modulated weakly (Misra and Pessoa, 2025).

This point is echoed by a growing literature. Simulated lesions showed that the effect of removing a region on network dynamics follows from its position in the connectome rather than from its activity (Alstott et al., 2009), and control-theoretic work ranks regions by their capacity to move the system rather than by how much they respond (Gu et al., 2015; Deco et al., 2019). Possibly, the sharpest form of the argument comes from neural populations, where activity counts only insofar as it projects onto relevant directions. Signaling between cortical areas is confined to a low-dimensional *communication subspace*, so much of a “source” area’s activity never reaches its “target” (Semedo et al., 2019), and preparatory activity is arranged to lie in an output-null subspace, large in amplitude and without effect on movement (Kaufman et al., 2014). The same type of geometry operates here at the level of regions. A region moves a state switch only through the part of its activity that points along the direction separating the two states, and whatever is orthogonal to that direction leaves the switch untouched (formally, a region contributes its activity weighted by *a*_*m*_, the projection of its emission loading onto the direction favoring state *j* over state *i*). What the present analysis adds is that the same region can occupy two roles within one model, steering a state and ending it, and that the two need not coincide. Amplitude maps remain a natural description of a region’s responds; they are not, on their own, a description of what it *does* to the system.

### Limitations

Several limitations of the present work deserved explicit discussion. The model is phenomenological rather than mechanistic: it establishes that the brain switches among distinct dynamical regimes but does not address what biophysical processes cause those regimes or drive the switches. Linearity within states is a simplification adopted for tractability, while the attractor landscapes suggested by theoretical work are considerably more complex (Miller, 2016; Rabinovich et al., 2006). Our piecewise linear approximation captures recurring patterns in the BOLD signal without modeling the nonlinear computations that generate them, a reasonable assumption given the hemodynamic timescale. The attractor interpretation is model-relative: stability is established within each learned linear regime, and our results should be read as identifying effective attractors at the fMRI timescale (Khona and Fiete, 2022). Markovian switching ignores longer histories, which semi-Markov extensions could capture (Oh et al., 2008; Shappell et al., 2019). A Markov chain forces geometrically distributed dwell times, and modeling the sojourn distribution directly changes the dwell times recovered from resting fMRI (Shappell et al., 2019). The recovered repertoire depends on the number of states and latent dimensions; other choices could subdivide or merge the states reported here. Subcortical claims from this model also carry a caveat of scale: the 54 subcortical regions are 21% of the parcellation but carry 3.5% of the emission loading, so the latent space couples to them through a small fraction of its total weight. The striatal and thalamic steering of State 1 should be read against that baseline. Global signal regression was applied during preprocessing and can influence anticorrelations and dynamic estimates (Saad et al., 2012; Chai et al., 2012). Resting fMRI is sensitive to physiological and vigilance fluctuations, and simultaneous physiology, pupillometry or EEG would help dissociate neural from non-neural contributions (Tagliazucchi and Laufs, 2014; Wang et al., 2016).

## Conclusions

Resting-state dynamics is neither a sequence of clean toggles nor a single smooth trajectory. Some transitions between whole-brain states are sharp at the limit of what fMRI can resolve, with the estimated flow reversing between adjacent samples (720 ms); others are gradual and one-sided. The heterogeneity is systematic, is unlikely to be attributable to hemodynamic filtering or to the discrete architecture of the model, and is recovered by analyses with different degrees of dependence on the model. Transitions have identifiable regional drivers whose influence accumulates before the switch and dissipates after it, and those drivers are not necessarily the regions that steer dynamics within the states involved. Treating state transitions as measurable dynamical objects, rather than as changes in a label, brings a set of questions about resting-state brain dynamics within empirical reach.

## Acknowledgments

L.P. is supported by the National Institute of Mental Health (R01 MH071589). Data for this study were provided in part by the Human Connectome Project, WU-Minn Consortium (Principal Investigators: David Van Essen and Kamil Ugurbil; 1U54MH091657) funded by the 16 NIH Institutes and Centers that support the NIH Blueprint for Neuroscience Research; and by the McDonnell Center for Systems Neuroscience at Washington University. The funders had no role in study design, data collection and analysis, decision to publish, or preparation of the manuscript.

## Methods

### Data

We employed resting-state fMRI data from the Human Connectome Project 1200-subjects release, processed with the ICA-FIX pipeline (Smith et al., 2013). For each resting-state fMRI run we first normalized (z-scored) the ROI time series, and then regressed out the global signal (the mean across all ROIs at each time point) from each ROI time series (but see supplementary analyses). Motion correction was performed voxelwise by regressing out 12 nuisance variables (six motion parameters and their temporal derivatives) using AFNI’s `3dDeconvolve` and `3dREMLfit` programs (Cox, 1996). The repetition time was 720 ms. HCP participants with 4 runs were considered. A set of *n* = 100 participants was used for exploratory analysis and model selection, that is, to select the number of SLDS states (*K* = 6) and the number of latent dimensions (*D* = 10); some supplementary analyses were also performed with models with four and eight states. A model was then fit to a separate set of *n* = 500 participants; all results reported here come from this model. We did not use all available HCP data because of the computational demands of estimating more than 500 participants; preliminary assessment indicated that SLDS model fits generated similar results after a sample size of approximately 200 participants.

#### ROI parcellation

We used a 254-ROI whole-brain parcellation, combining the Schaefer–Yeo 200-ROI cortical parcellation (Schaefer et al., 2017) and the Tian 54-ROI subcortical parcellation (Tian et al., 2020). Canonical networks were based on the Yeo-17 parcellation (Yeo et al., 2011).

### Model

A multi-level SLDS model was fit to the HCP resting-state data, following the general proposal of Linderman et al. (2019): the multi-level formulation allows one to simultaneously estimate both individual and group-level dynamics, enabling quantification of the variability of individual-level parameter estimates. The model comprises an SLDS component and a multi-level component.

### SLDS component

The SLDS model includes three parts: dynamics, transition, and emission. A set of brain states is inferred from the multivariate time series data 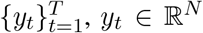 (here *N* = 254 ROI time series). Each state is associated with its own linear *dynamics*. The probability of a state *transition* at any time point depends on both the current state label (*z*_*t*_ *∈* {1, *…, K*}) and the system’s current position in the continuous latent state space (*x*_*t*_ *∈* ℝ^*D*^). The *emission* model maps dynamics in latent space (ℝ^*D*^) to activity in ROI space (ℝ^*N*^), allowing model predictions to be visualized in the space of the brain.

- **Latent dynamics**. For each state *k*, latent dynamics is modeled as a linear equation

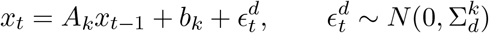

where *A*_*k*_ *∈* ℝ^*D×D*^ is the state dynamics matrix, *b*_*k*_ *∈* ℝ^*D*^ is the state bias vector, and 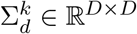 is the residual covariance matrix. Here *D* = 10.
- **Transition between states**. For any two states *i, j*, the transition probability is modeled as

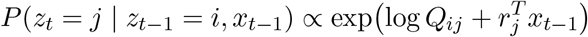

where *Q ∈* ℝ^*K×K*^ is the Markovian transition matrix and *r*_*j*_ *∈* ℝ^*D*^ is the state recurrent coefficient acting on the latent continuous state *x*. The vector *r*_*j*_ acts as a set of “weights” for state *j*: a strong match between the current latent position *x*_*t−*1_ and *r*_*j*_ increases the probability of transitioning into state *j*. This mechanism allows the observed brain dynamics to directly influence and bias the latent state transitions, moving beyond purely fixed Markovian probabilities.
- **Emission model**. The emission model links fMRI observations *y*_*t*_ *∈* ℝ^*N*^ and latent state *x*_*t*_ *∈* ℝ^*D*^ through a linear model:

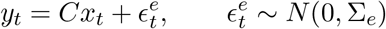

where *C ∈* ℝ^*N ×D*^ is the emission matrix and Σ_*e*_ *∈* ℝ^*N ×N*^ is diagonal. The emission matrix is assumed unchanged across states.

### Multi-level component

We assumed that each participant has their own dynamical system (i.e., parameters), and employed a multi-level Bayesian framework to estimate a group-level model while respecting participant-level variability. Each subject was associated with their own participant-level SLDS parameters 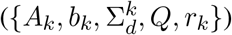 linked hierarchically to group-level SLDS parameters. Emission parameters (*C* and Σ_*e*_) were assumed unchanged across subjects given that they do not model the underlying brain activity.

1. **Hierarchical prior for intrinsic system parameters**. Let *θ* denote the collection of all parameters of the dynamics and transition elements of an SLDS model:

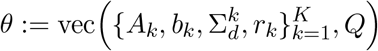

where vec stands for vectorizing. For a total of *n* subjects, we link the subject-level parameters to group-level ones by a hierarchical prior (using superscript *i* to denote subject index)

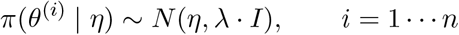

where *η* denotes the group-level parameters corresponding to *θ, I* is the identity matrix, and *λ* determines how strongly subject-level parameters are linked to the group parameters (we used *λ* = 0.1). The identity covariance prior indicates that no prior information is placed on the covariance structure among parameters. The hierarchical posterior *f* (*θ* | *η*, **y**) is inferred based on fMRI data. Values of *λ* in the range 0.01 to 0.5 yielded qualitatively similar group-level SLDS results.
2. **Emission parameters are shared across subjects**. As the emission part of SLDS is more linked to mapping dynamics back to the brain space, and less about the intrinsic system being modeled (in latent space), we assumed that the emission parameters do not vary across subjects.

All group and subject level parameters were fitted jointly, yielding one group-level model and *n* subject-level models. The optimization procedure adapts the variational Laplace EM algorithm (Zoltowski et al., 2020) to support hierarchical Bayesian parameter fitting; for a related approach, see Linderman et al. (2019). Specifically, the EM procedure for SLDS (Zoltowski et al., 2020) was extended to account for the fMRI data structure: in the E-step (state update), the variational Laplace approximation was carried out individually using the current subject-level SLDS parameters to update latent states 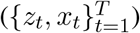 per subject; in the M-step (parameter update), group-level and all subject-level SLDS parameters 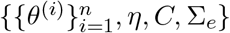 were updated simultaneously. See Appendix A for details.

### Parameter selection

A separate set of 100 participants was used to determine two key parameters of the model via 10-fold cross validation.

#### Number of states

The information-theoretic measures Adjusted Rand Index and Normalized Mutual Information were used to quantify state *consistency* (i.e., the sequence of labels estimated by the model) among cross-validation fits, and to choose the number of states *K* (Vinh et al., 2010), similar to our previous paper for a task paradigm (Misra and Pessoa, 2025). Based on the consistency values at *K* = 6 and their reasonable variability among different latent dimensions, we chose six states.

#### Number of latent dimensions

To select the dimensionality of the latent space, we used an approach based on explained variance (one-step prediction accuracy of the dynamic model, under noiseless dynamics). Motivated by the *R*^2^-like measure previously defined for a linear dynamical system (Liégeois et al., 2019), for the SLDS case let

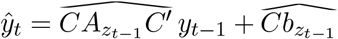

be the one-time-step model prediction of *y*_*t*_ based on the previous state (*C*^*′*^ denotes the transpose of *C*). The prediction residual is 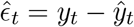, and the proportion of variance explained by the SLDS model is

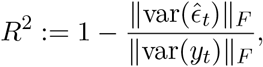

where ∥ · ∥_*F*_ denotes the Frobenius norm. The increase in explained variance slowed down around *D* = 10, while adding dimensions would considerably increase both memory and computational complexity (*O*(*D*^2^)). Explained variance varied little with the number of states *K*.

### Definitions and statistical approach

#### State activity maps

Activity maps display, for each state, mean activity at “equilibrium” time, defined as the state mean duration plus 3 samples.

#### Network activity

Networks were based on the Yeo-17 parcellation. Activity was defined by averaging the activity values of all ROIs within a given network.

#### Temporal activity during transitio

Brain activity was averaged across all state transition segments; *t* = 0 denotes the transition time point, that is, the first sample of the new state.

#### Statistical approach

We sought to perform statistical tests by considering individual participant-based variability. For results at the map level, individual-level maps were determined followed by a one- or two-sample *t*-test across participants. To control for multiple comparisons, we used a false discovery rate approach such that the expected proportion of false positives was controlled at 5% (Benjamini and Hochberg, 1995). Additional tests were also based on individual-level values; where appropriate, Bonferroni correction was applied based on the specific number of tests.

#### Testing stability of state dynamics

Linear system dynamics is stable when all eigenvalues of the dynamics matrix lie within the unit circle (Luenberger, 1979). For each state, we tested for stability at the group level by using subject-level model values of the maximum eigenvalue norm (spectral radius) of the state dynamics matrix. For state *k*, let *ρ*_*k*_ := max{|*λ*| : *λ ∈ σ*(*A*_*k*_)} (*σ* denotes the set of eigenvalues), and 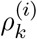 denote the *ρ*_*k*_ value from the subject-level SLDS for subject *i*. Our test statistic was defined as 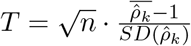, where 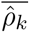 and *SD* 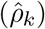 are the sample mean and standard deviation of fitted subject-level values. In other words, 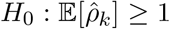 versus 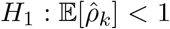. All state dynamics were stable at the group level (*p* < 0.05, Bonferroni correction for *K* = 6 states).

### Transition matrix and statistical significance

Transition probabilities were computed from state label sequences at the individual level and averaged. To quantify state *transitions*, we were interested in occurrences of state *i* at time *t* followed by state *j* ≠ *i* at time *t* + 1 (i.e., we were not interested in self-transitions). Entries {*i, j*}, *i* ≠ *j*, indicate the probability of transitioning from state *i* to state *j*.

Statistical significance of state transitions was determined by a group permutation test (*B* = 1000 permutations) as follows. For *b* = 1, · · ·, *B*: (*Step 1*) for every subject *s*, a randomized state label time series was generated by permuting the states by considering same-state segments (e.g., 114455 could be shuffled to 441155, 445511, etc.); (*Step 2*) for every subject *s* and transition *i* ↦ *j* (*j* ≠ *i*), compute the transition probability from the randomized state label time series, denoted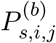; (*Step 3*) denoting the observed transition probability by 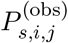, compute for every subject and transition a z-score 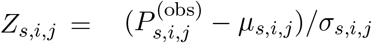 where *µ*_*s,i,j*_ and *σ*_*s,i,j*_ are the sample mean and standard deviation of the permuted probabilities; (*Step 4*) for every transition, perform a one-sample *t*-test on 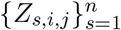 to test if they are greater than zero. Significance was determined at *p* < 0.05 after Bonferroni correction for (*K*−1) × *K* state transitions.

### Region dynamics importance

Following our prior work (Misra and Pessoa, 2025) we computed region dynamics importance. The SLDS model can be used to predict how fMRI brain signals evolve in one time step:

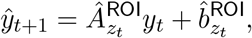

where 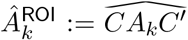 denotes the estimated dynamics matrix for state *k* in brain space ℝ^*N*^, and 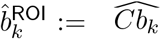 denotes the estimated bias term in the same space.

We define the importance of the *i*th brain region at time *t* by evaluating the whole-brain activity difference at *t* + 1 when zeroing out (“lesioning”) the *i*th column in *A*^ROI^ (the resulting matrix is denoted by *A*^ROI^[:, −*i*]):

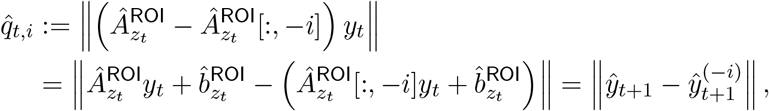

where *∥* · *∥* denotes the Euclidean norm in ℝ^*N*^, and 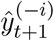 provides the estimated activity supposing region *i* is zeroed. Computing an importance value at each time point yields an “importance time series”, which was normalized (z-scored) for each run. Note that

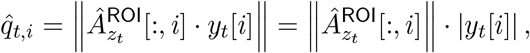

so that the measure takes into account both the activity of region *i* itself and the “effective connectivity” from region *i* to other brain regions.

#### Statistical test

To threshold importance maps we tested against the null hypothesis that all regions have equal importance, by shuffling ROI labels. Let *Q*_*s,k,i*_ denote region *i*’s importance in state *k* for subject *s*. For every state *k*, region *i*, and subject *s*: (*Step 1*) compute observed importance values 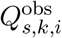 (*Step 2*) for *b* = 1, · · ·, *B* (*B* = 1000), generate a permutation 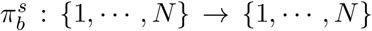 (*Step 3*) z-score the observed importance value based on the range of values obtained by the permuted (“null”) values, 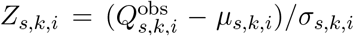 Then (*Step 4*) test if the z-scored importance is greater than zero via a one-sample *t*-test on 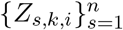. Multiple comparison correction was applied for *K* states and *N* regions via false discovery rate control at 5%.

### Region transition importance

The SLDS model estimates transition probabilities, which can be exploited to investigate the contribution of individual brain regions to state transitions. Recall that SLDS models a state transition *i* ↦ *j* as *P* (*z*_*t*+1_ = *j* | *z*_*t*_ = *i, x*_*t*_) *∝* exp 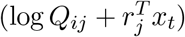. For a given state *i* we can define *ĥ*_*i*_(·) : ℝ^*D*^ → ℝ^*K*^ :

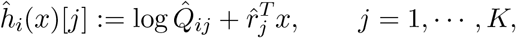

for a given system position *x* in latent space. Our goal is to quantify the importance of region *m* for state transition *i* ↦ *j*.

**Step 1: Computing the latent position via leave-one-region-out projection**. At a given time *t*, an estimate of the system’s position in latent space is obtained via the emission model. Because the fitted emission matrix has orthonormal columns ( *∥ Ĉ*^*⊤*^*Ĉ* −*I ∥*_max_ = 1.1 × 10^*−*15^ for the model reported here), that estimate is the projection 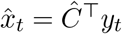, in which each region contributes an additive term *ĉ*_*m*_ *y*_*t*_[*m*] with *ĉ*_*m*_ = *Ĉ* [*m*, :]^*⊤*^. Lesion of region *m* is done by removing that region’s term,

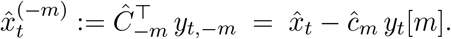

The goal is to isolate how much information ROI *m* uniquely contributes to the model’s tendency to leave state *i* for *j*. By dropping the term ROI *m* contributes, the method constructs a “counterfactual” latent 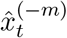 built from the rest of the brain *without letting ROI m influence the latent estimate*. We note that this is not identical to re-solving the least-squares problem over the remaining regions: for the full orthonormal *Ĉ* the two coincide, but once a row is removed the reduced matrix is no longer orthonormal and the two estimates differ. The procedure used here, and in all results reported below, is the projection above.

**Step 2: Computing the log-odds of the state transition probability**. At a given time *t* we measure the change in state transition probabilities resulting from lesion of region *m* by computing an instantaneous log-odds in favor of the transition *i* ↦ *j* versus staying in state *i*:

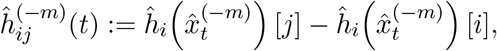

which turns the lesioned latent estimate into a decision-relevant quantity: the instantaneous log-odds that the system leaves state *i* for *j* rather than remaining in *i*. Because the normalizing constant of the transition model is shared by all destinations, it cancels in this difference, so the quantity is an exact log-odds rather than an approximation (measured in nats and its exponential is an odds ratio). Using the difference puts the effect of ROI *m* on a *relative* scale, isolating how the lesioned evidence affects the transition versus stay option; higher values thus indicate a stronger immediate drive toward *j* under the lesion of region *m*.

**Step 3: Averaging over all state transitions** *i*↦ *j* **observed at time** *t*. Let *T*_*c*_ be the collection of all instances of state transitions *i* ↦ *j* observed in the time series at time *t*. We average these instances to compute

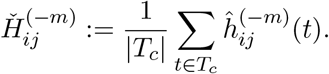

#### Definition

We define *region transition importance* for region *m* as

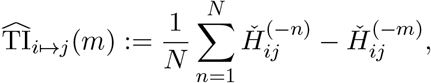

where the first term is the average across all ROIs. A positive TI value means that region *m* has above-average contribution for *i* ↦ *j*, such that lesion of region *m* tends to reduce transition probability.

Substituting Step 1 into Steps 2 and 3 gives a closed form. Writing 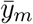 for the mean activity of region *m* over the pre-transition window and instances,

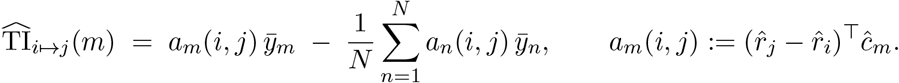

Transition importance is therefore the pre-transition activity of each region, weighted by a purely geometric coefficient *a*_*m*_ that measures how far that region’s emission loading points along the direction favoring *j* over *i*, and then centered across regions. It is noteworthy that log cancels identically, so the 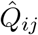 measure carries no information from the baseline transition matrix.

#### Transition importance map

Transition importance maps were generated via a procedure similar to Step 3. Instead of considering values 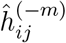 at just one time point *t*, we averaged time points across a window before the transition time: observations during three time steps, *t ∈* [−3, −1], before the transition *i* ↦ *j* (considering other windows, such as *t ∈* [−5, 0], did not qualitatively change the result). To threshold the maps statistically we used the same testing procedure as for dynamics importance maps, testing against the null of equal importance across all regions. Multiple comparison correction was applied taking into account the number of transitions times the number of regions.

#### The drive toward a switch

Region transition importance weights the activity of each region by *a*_*m*_(*i, j*), and then centers those terms across regions. The centering is what makes the measure a comparison between regions, and it removes their total. Here we keep the total. Each region contributes *a*_*m*_(*i, j*) *y*_*t*_[*m*]: its activity, weighted by how far its emission loading points along the direction that favors *j* over *i*. Summing the contributions of all *N* regions gives the drive toward the switch,

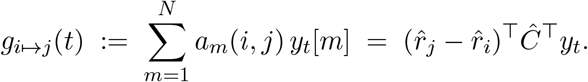

The two expressions are equivalent but one describes it region by region, the other as a single projection of the whole-brain pattern onto the direction that favors *j*. Regions with positive terms push the system toward the switch, and regions with negative terms hold it back.

The model gives the instantaneous log-odds of switching rather than staying as log 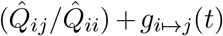. The first term is a constant of the fitted model, and the second moves with the brain; *g* is the second alone. A value of zero therefore does not mean that the switch is impossible. It means that the current pattern is neutral, and leaves the odds at whatever the baseline transition matrix assigns them. Because *g* is in *nats*, exp(*g*) is the factor by which the current pattern multiplies those baseline odds.

We evaluated *g* on *Ĉ*^*⊤*^*y*_*t*_, the projection of the measured signal at frame *t*, rather than on the posterior latent trajectory. This is the estimate that region transition importance is built on, which is what makes the regional terms sum to *g* exactly. Because *g* is a log-odds and not a probability, we referenced the curves of Fig. 6 to a mid-dwell level rather than to zero: for each participant and transition we evaluated *g* at the midpoints of dwells of at least fifteen frames in the departing state, and subtracted that value. Confidence intervals are bootstrap intervals over participants (*B* = 2,000), with the baseline subtracted inside each replicate. The regional shares of Fig. 7 are the same terms 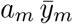, averaged over the window *t ∈* [−3, −1] and the same instances, divided by their sum.

### Quantifying transition dynamics via vector field contrast

Based on the state dynamics matrix *A*_*k*_, we define a state-specific vector field representing how a point in state space flows under model dynamics. For any point *x* in the *D*-dimensional latent space belonging to discrete state *z*, the SLDS model specifies a one-step change as

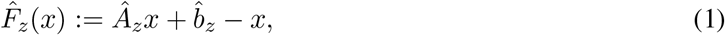

that is, the expected evolution of the latent trajectory *x*_*t*_ in one time step.

#### Field contrast

To measure how different the equations of motion of two states are, we define a measure based on the *L*_2_ distance between the vector fields of the two states, origin and destination. For transition *i* ↦ *j*, the expected difference between the two vector fields *F*_*i*_ and *F*_*j*_ (estimated by Eq. 1) is estimated by the average Euclidean distance between vectors over sample points,

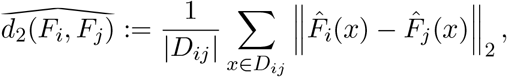

where *D*_*ij*_ stands for a set of points *x* combining all state *i* points preceding state transition and all state *j* points post transition. We take 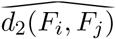 I\as the *field contrast* of the transition, computed per subject and per transition of interest. The quantity is a distance between two rules of motion evaluated where the system actually is; it says how much of the expected one-step displacement is altered when one rule replaces the other, and it says nothing about how long that replacement takes, which in an SLDS is one time step by construction. Being a distance between vectors, it also does not separate a change in the direction of the expected displacement from a change in its length.

The field contrast is defined in the full ten-dimensional latent space and was computed for the eight transitions reported here. The two transitions involving State 6 are excluded throughout: a state whose mean dwell is 2.9 s does not supply the eleven consecutive within-state frames the mid-dwell floor requires. We note that State 2 ↦ State 1 and the two State 6 transitions all showed weaker fMRI activity changes, smooth across most canonical networks (Fig. S3).

#### Vector field as a function of time

Equation 1 defines a single vector field for a state. To describe the finer temporal evolution of vector fields, we consider a one-step difference:

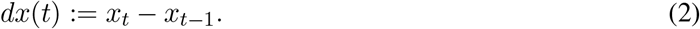

A one-step difference spans two frames and can be named after either of them; we name it after the frame it *arrives* at. Writing *τ* for the first frame carrying the new label, the displacement that crosses the boundary, *x*_*τ*_ − *x*_*τ−*1_, is therefore *dx*(*τ*) and is assigned to time zero. Two considerations settle this choice. The switching model governs the step from *x*_*t−*1_ to *x*_*t*_ by the state occupied at *t*, so the arriving frame is the one whose dynamics generate the step; and with this convention the time axis of the fields coincides with that of the activity analyses above, in which *t* = 0 is the first sample of the new state. The alternative convention, naming a displacement after the frame it leaves, shifts every lag reported here by one and places the boundary-crossing step at *s* = −1.

To estimate the vector field at time *s* for transition *i* ↦ *j*, consider a local area ℬ in the state space centered at point **a**. We compute an average 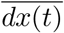 over all times positioned at time *s* during an *i* ↦ *j* transition period, and for which the latent trajectory *x*_*t*_ enters the local area ℬ:

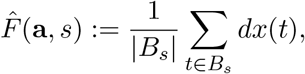

where *B*_*s*_ := { *t* : *t* = *τ* + *s* for some *i* ↦ *j* transition point *τ, x*_*t*_ *∈* ℬ}. Note that this field is estimated from the inferred latent trajectory rather than from the state dynamics matrices, and is therefore free to vary continuously in time.

#### 2D visualization of vector fields

To visualize 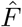 (**a**, *s*) in two dimensions, we sample points uniformly from a grid on the plane spanned by the first two latent dimensions determined by SLDS, which are of largest variance as well as of largest one-step predictive variance (*R*^2^). Let *x*_2*D*_ denote the projection of a *D*-dimensional vector *x* onto this plane, and *F*_2*D*_ denote the vector field after projection. For any grid point **a** = (*a*_1_, *a*_2_), we collect time points for which the 2D projection of the latent trajectory (*x*_*t*_)_2*D*_ enters a rectangular cell ℛ:= [*a*_1_ − *δ*_1_, *a*_1_ + *δ*_1_] × [*a*_2_ − *δ*_2_, *a*_2_ + *δ*_2_] (where *δ*_1_ and *δ*_2_ determine the grid resolution) centered at **a**. The inferred 2D vector at **a** is determined by the average *dx*(*t*) over the collected time points:

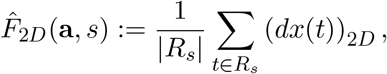

where *R*_*s*_ := { *t* : *t* = *τ* + *s* for some *i* ↦ *j* transition point *τ*, (*x*_*t*_)_2*D*_ *∈* ℛ}, and *dx*(*t*) is defined in Eq. 2. The notation (*dx*(*t*))_2*D*_ means that the one-step difference *dx*(*t*) is first computed in the *D*-dimensional state space and then projected onto the 2D plane.

#### Cosine similarity between adjacent fields

To characterize how quickly the flow reorients, we computed the cosine similarity between 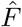(**a**, *s*) and 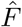 (**a**, *s* − 1), that is, between the field at lag *s* and the field one time step earlier. This is the transition sharpness index. It was computed in the full *D* = 10 latent space. The local areas ℬ were defined by *k*-means on the latent positions, using 121 clusters, the same number of cells as the 11 × 11 grid of the two-dimensional visualization above. The clustering was fitted once per transition, on that transition’s own pooled positions across all lags, and then held fixed: it defines the estimator’s neighborhoods, and is not re-estimated when the data are resampled. The field at lag *s* in neighborhood **a** is the mean of the ten-dimensional *dx*(*t*) over the points of that neighborhood. A neighborhood entered the average at lag *s* only if it held at least ten points at both *s* and *s* − 1; sparse neighborhoods were dropped rather than imputed.

#### A floor for the cosine

A cosine between two averages estimated from finite samples is attenuated toward zero, by an amount that depends on how many points contribute and how noisy they are. A value near zero is therefore not by itself evidence that the flow reversed. We measured the size of that attenuation directly rather than modeling it. For each transition *i* ↦ *j* we assembled a set of *pseudo-transitions*: time points in the middle of a dwell, with the state label constant over [*τ* − 5, *τ* + 5], drawn from both state *i* and state *j*. No change of rule occurs at such a point, so whatever the estimator returns there is attenuation and noise alone. The pseudo-transitions were then passed through the identical estimator, subsampled at random so that every (lag, neighborhood) pair carried exactly the same number of contributing points as the real data. The null therefore differs from the data in one respect only, namely that no transition occurs. We report both the observed cosine and this floor, and the excess of one over the other.

#### Confidence intervals and the criterion for a reversal

Intervals were obtained by subsampling participants: 200 replicates, each drawing *m* = 250 of the 500 participants without replacement and recomputing both the observed cosine and its count-matched floor on that subset, with the standard deviation across replicates scaled by 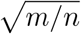 to give the standard error at the full sample size. We used subsampling rather than a with-replacement bootstrap because duplicating participants reduces the number of *distinct* points contributing to each neighborhood, which attenuates a cosine of two noisy averages in exactly the way described above; a percentile interval computed that way is displaced toward zero and can fail to contain the full-sample estimate.

A transition was counted as reversing the flow only when the 95% interval (*ĉ* ± 1.96 SE) at the crossing step lay entirely below zero, and only when it also survived correction for the number of transitions examined. For the latter we treated the eight transitions as eight one-sided tests of *H*_0_ : *c* ≥ 0 with *z* = *ĉ/*SE, for which Bonferroni correction gives *α* = 0.05*/*8 = 0.006 and a threshold of |*z*| = 2.50. All four reversals cleared it, the weakest at |*z*| = 3.9. Because the criterion is applied at a single lag fixed in advance, no further correction across lags is required, and the transitions are reported individually rather than pooled. We note that the standard errors are themselves estimated from 200 replicates, so the larger *z* values should be read as indicating that a reversal is far from the threshold rather than as exact tail probabilities.

### State transition boundaries

Two kinds of quantity are plotted against a transition in this paper, and they mark the boundary at different places, as clarified here. A quantity defined at a single frame (network activity, the drive toward a switch, and region transition importance) is plotted against the frame index relative to the boundary. The boundary itself lies *between* two frames, so the last sample of the departing state is −1 and the first sample of the new state is 0, and a quantity that accumulates while the system is still in the departing state necessarily reaches its maximum at −1 (Figs. 6 and 8). A quantity defined on the step between two frames (the vector field and the cosine between adjacent fields) has no single frame to stand on, and we name each displacement by the frame it arrives at. The step that crosses the boundary runs from the last sample of the old state to the first sample of the new one, and is therefore labeled 0 (Figs. 4 and 5; the choice is justified in “Vector field as a function of time” below). The two conventions agree because the crossing step is precisely the step that arrives at frame 0: an accumulating quantity peaking at −1 and a flow reversing at 0 describe the same instant, the passage from the last sample of one state to the first sample of the next. Finally, note that nothing in this paper is resolved more finely than that single 720 ms step.

### Do the regions that steer a state also end it?

#### Destination-averaged control

Transition importance is defined for a directed transition, whereas dynamics importance is defined for a state, and three of the origin states have two significant exits. A mismatch between the two rankings could therefore reflect the destination-specificity of transition importance rather than any difference between steering a state and ending it. To test this, we collapsed each origin state’s exits into a single map, both as an unweighted mean over its transitions and as a mean weighted by the number of instances of each, and repeated the comparison against that state’s dynamics importance. State 2 and State 4 have one significant exit each, so for them the averaged map is the original one. The result did not change. State 1 shared none of its top ten and none of its top twenty with the averaged map, its steering regions ranking 201st to 254th of 254 for ending it; State 5 also shared none, ranking 25th to 125th; and State 3 shared four of ten and eleven of twenty, ranking 1st to 131st. We also compared the transition importance maps with each other. Two exits from the same state correlate at *r* = 0.58 on average, barely above the *r* = 0.44 of transitions that share no state at all, whereas the two directions of the same state pair correlate at *r* = 0.94. Transition importance is therefore carried mainly by the pre-transition activity pattern, which the two directions of a pair share and which differs little between a state’s exits, so the comparison reported here does not depend on which destination is considered.

## Supplementary figures

**Figure S1:**
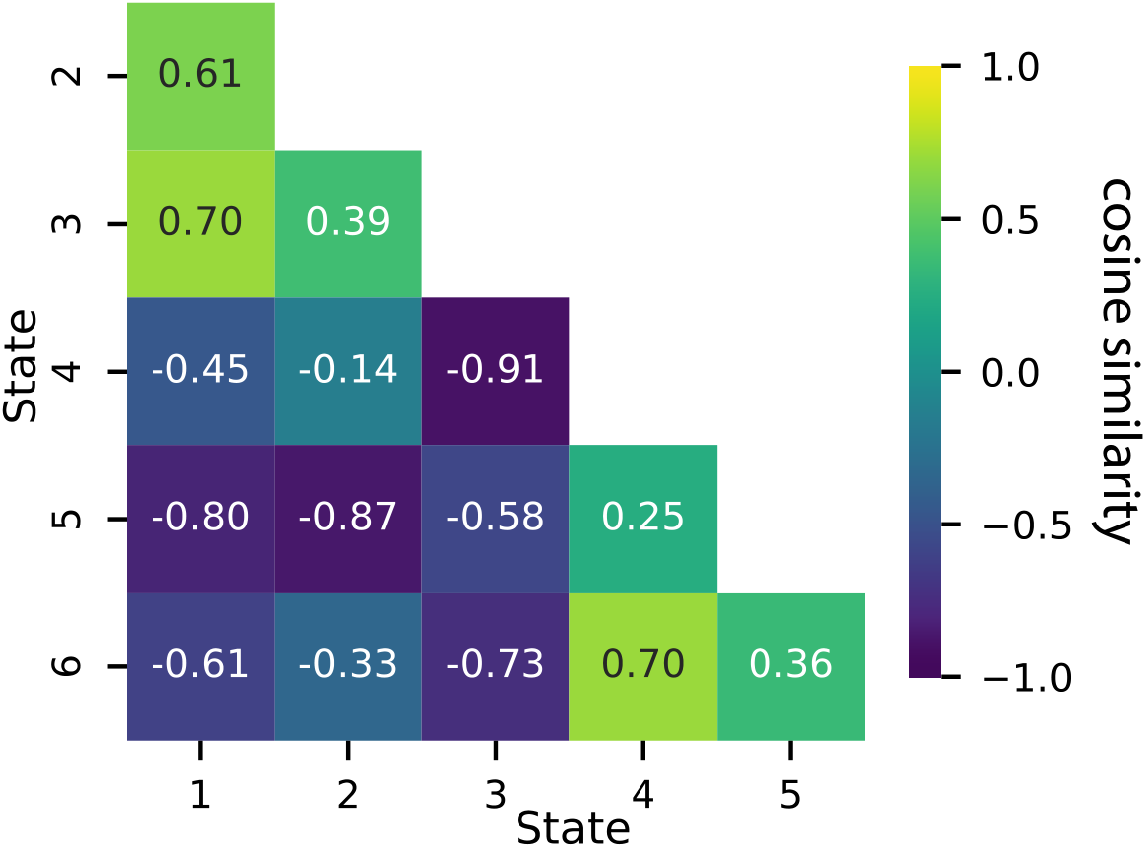
State similarity of fMRI activity. Activity maps at “equilibrium” were compared via cosine similarity (both cortical and subcortical regions were considered).

**Figure S2:**
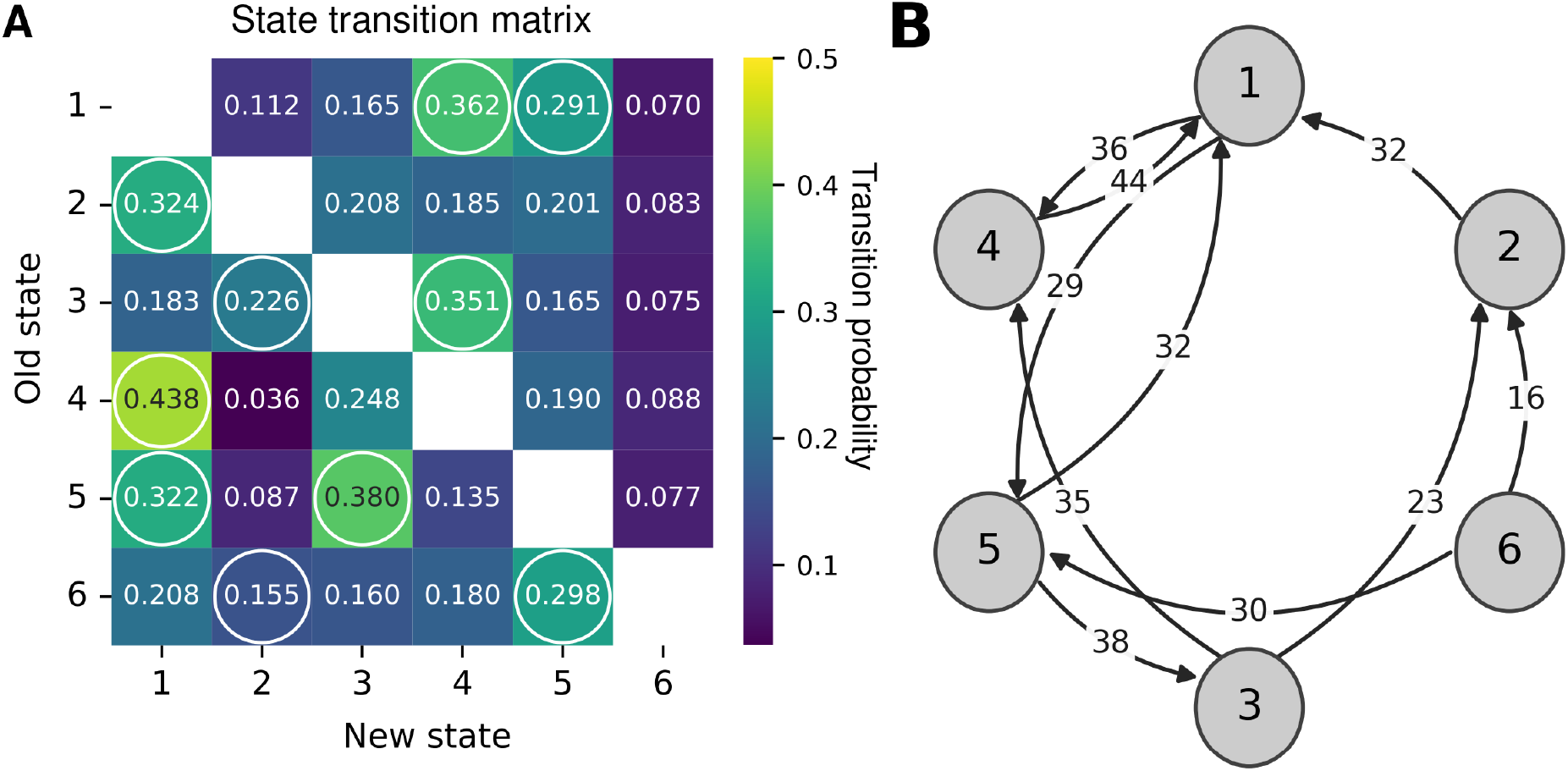
State transition probabilities. **A**. State transition probabilities (excluding self-transitions). Circled entries indicate statistically significant transitions (permutation test, *p* < 0.05, multiple comparison corrected for 6 × (6 −1) transitions). The diagonal is omitted as we did not consider self-transitions. **B**. Transition graph for significant state transitions.

**Figure S3:**
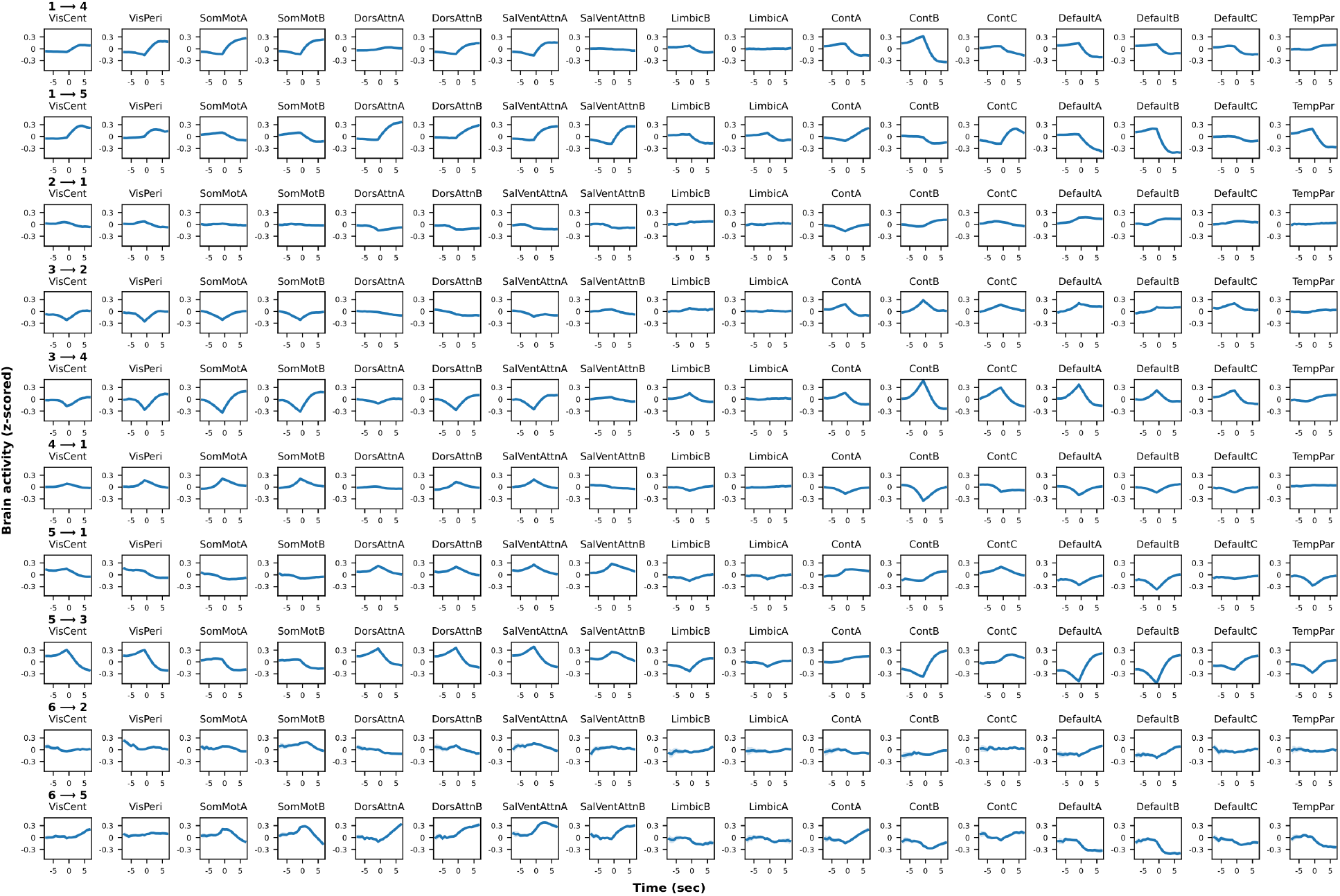
Temporal activity for each cortical network during significant state transitions. *t* = 0 indicates the first time point of the new state. Transitions involving State 6 are shown here for illustration only; they were not considered in the paper because the state’s mean dwell time was too brief to allow meaningful quantification of features of state transitions.

**Figure S4:**
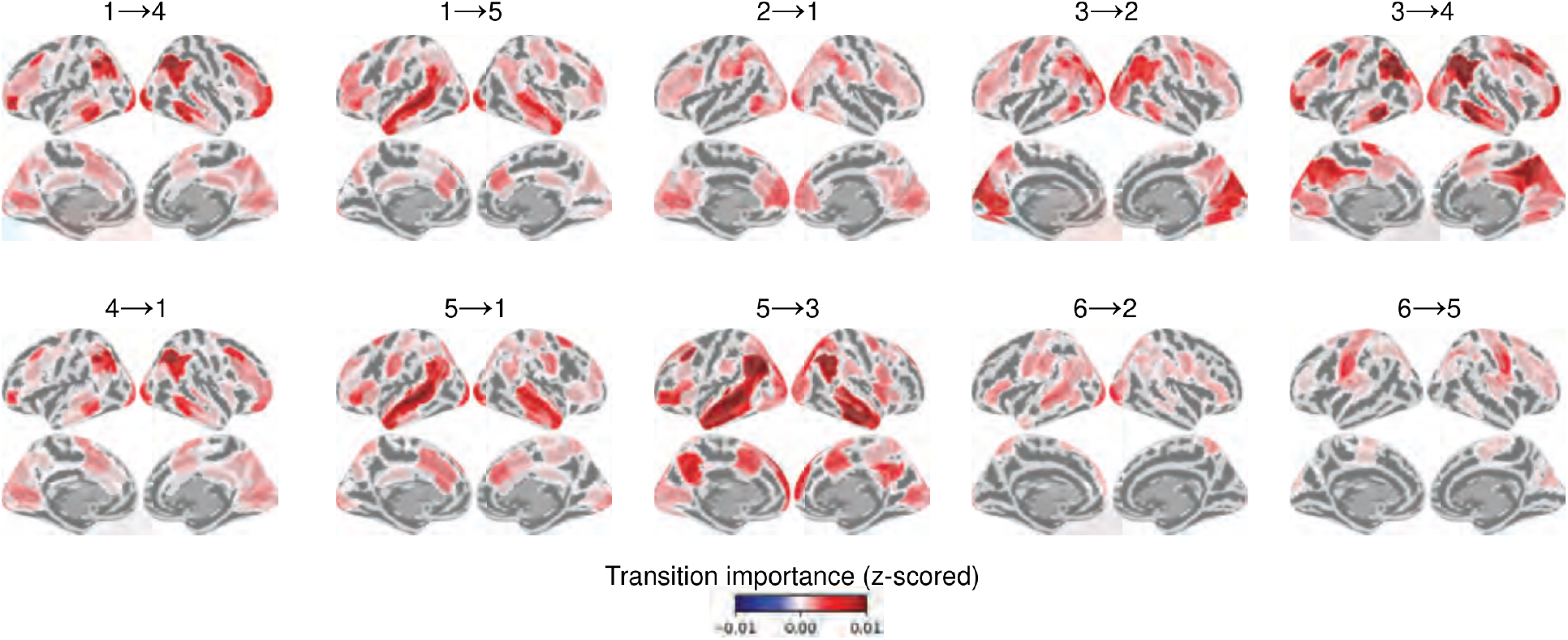
Transition importance maps for significant transitions. Maps are thresholded against the null of equal importance across regions (False Discovery Rate at 0.05). Transitions involving State 6 are shown here for illustration only; they were not considered in the paper because the state’s mean dwell time was too brief to allow meaningful quantification of features of state transitions.

**Figure S5:**
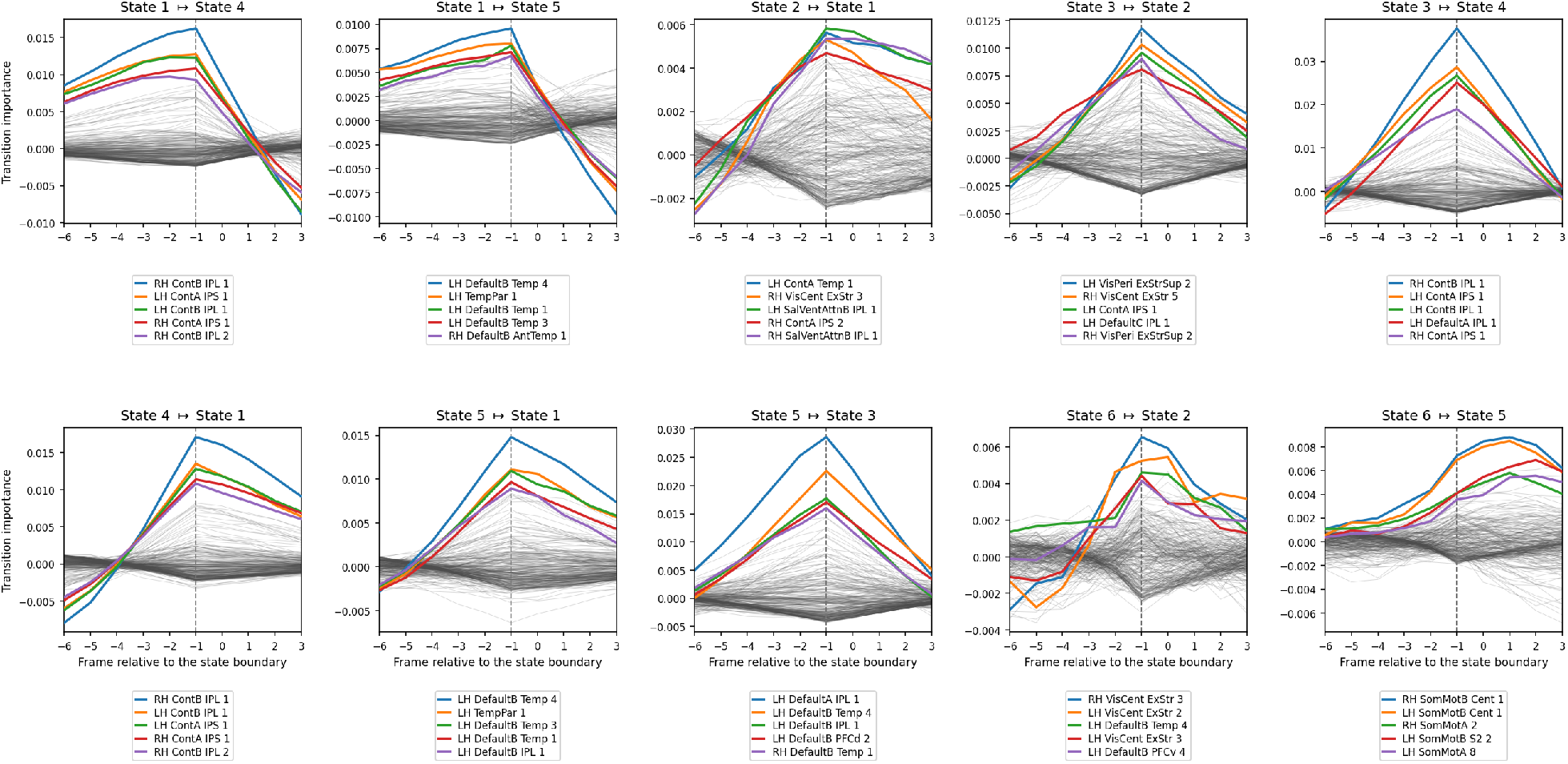
Temporal change of transition importance values during significant state transitions. Each curve represents a brain region, with the top five highlighted. The axis is in frames relative to the state boundary, as in Figs. 6 and 8: frame 0 is the first sample of the new state and the dashed line marks frame −1, the last sample of the departing state. Transitions involving State 6 are shown here for illustration only; they were not considered in the paper because the state’s mean dwell time was too brief to allow meaningful quantification of features of state transitions.

**Figure S6:**
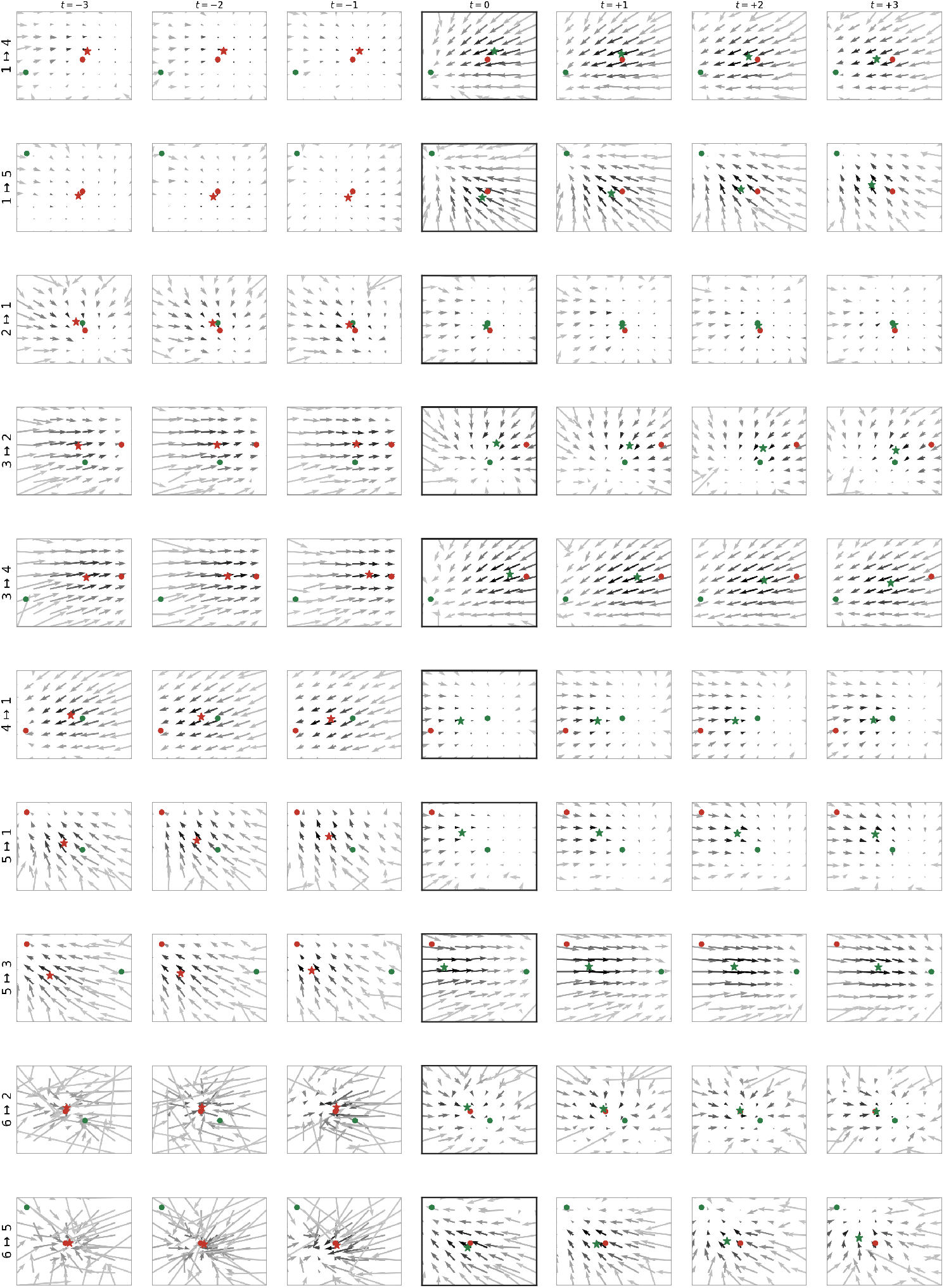
Vector field changes during statistically significant state transitions. Each panel is one time step. As in Fig. 4, a displacement is labeled by the frame it arrives at, so *t* = 0 (boxed) is the step that crosses the boundary, from the last sample of the old state to the first sample of the new one. Red and green dots are the attractors of the departing and arriving states, and the star is the mean position of the trajectory, red before the boundary and green from the crossing step onward. The grayscale of each vector is proportional to the number of data points falling in that cell. Transitions involving State 6 are shown here for illustration only; they were not considered in the paper because the state’s mean dwell time was too brief to allow meaningful quantification of features of state transitions. The fields of State 6 ↦ State 2 and State 6 ↦ State 5 are visibly less organized than the rest.

**Table S1:**
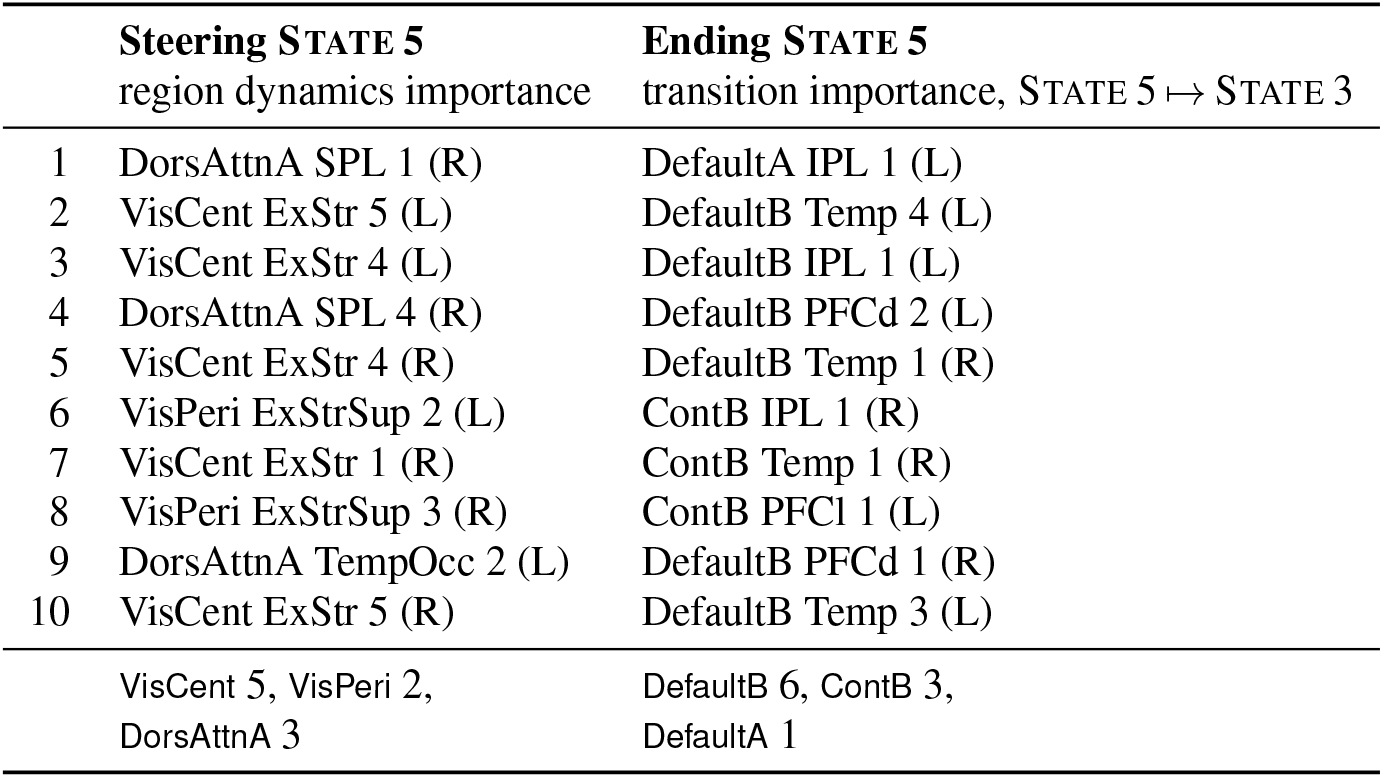
Regions that steer a state are not the regions that end it. The ten highest-ranked regions of State 5 by within-state region dynamics importance, which measures how much the predicted one-step evolution of whole-brain activity changes when a region’s contribution is removed from that state’s dynamics matrix (Misra and Pessoa, 2025), beside the ten highest-ranked by transition importance for State 5 ↦ State 3. Both rankings are averages over the 500 participants. The two lists do not share a single region: State 5 is steered by dorsal attention and visual regions, consistent with its attention/control character, and is ended by default-mode and control regions. The dissociation is not merely a reshuffling of the same set; the ten regions that steer State 5 rank between 24th and 77th of 254 for ending it. Labels are Schaefer–Yeo17 parcels with hemisphere in parentheses; the final row gives the network composition of each column.

## Appendix A: Optimization of multi-level SLDS

We outline the optimization framework for a two-level (hierarchical) nonlinear Bayesian model, with a focus on switching linear dynamical systems (SLDS). We begin by reviewing the general setup of a hierarchical Bayesian model and its MAP estimation. We then describe the Expectation–Maximization (EM) algorithm for a single SLDS and extend it to the multi-level case, showing how to jointly estimate both subject-level and group-level parameters.

### Hierarchical Bayesian model fitting for group studies

We consider a two-level model with subject-level (child) parameters *θ* and a group-level (parent) parameter *η*, along with observed data *y*. The key notations are listed in Table S2.

**Table S2:** List of notations in a generic hierarchical model.

|  |  |
| --- | --- |
| $\theta$ | single-level parameters or subject-level parameters (children) |
| $\eta$ | group-level parameters (parent) |
| $y$ | observations (e.g., emissions in an SLDS) |
| $f(\cdot)$ | likelihood function |
| $l(\cdot)$ | log-likelihood function |
| $\pi(\cdot)$ | prior probability density |

A hierarchical prior ties each subject’s parameter *θ* to the group parameter *η*. For example, we might assume

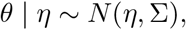

so that *π*(*θ* | *η*) is normal with mean *η* and covariance Σ.

In a non-hierarchical (single-level) parametric model, one would fit *θ* by maximum a posteriori estimation (MAP):

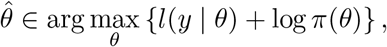

where *l*(*y* | *θ*) is the log-likelihood of the observations and *π*(*θ*) is the prior.

In the hierarchical model, we aim to estimate both *θ* and *η* from the data. The joint posterior density of (*θ, η*) given observations *y* is

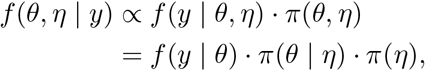

whose logarithm is

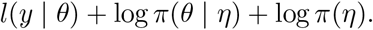

The term *l*(*y* |*θ*) is the same log-likelihood as before, while *π*(*θ* |*η*) is the additional term in the hierarchical model.

A straightforward approach is simultaneous MAP estimation of both levels, i.e., optimize child and parent parameters together:

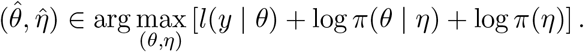

This yields estimates of both subject-level (*θ*) and group-level parameters (*η*). One caveat is that the total number of parameters grows linearly here with the number of subjects, but in practice modern optimizers such as SGD and ADAM can handle these large problems.

### EM for single-level SLDS

In a non-hierarchical (single-level) SLDS, let *θ* denote the set of SLDS parameters and (*x, z*) be the set of the latent continuous and discrete states. The EM procedure for SLDS (Zoltowski et al., 2020) proceeds as follows in the *t*-th iteration:

- **E-step**: Compute the expected complete-data log-likelihood (Q-function, Dempster et al., 1977) given the current parameter estimate *θ*^(*t*)^:

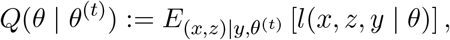

where expectation is with respect to the posterior of the latent states (*x, z*) under the current parameters (in practice this posterior is approximated by variational inference and Laplace approximation, we omit those details, which can be found in Zoltowski et al., 2020).
- **M-step**: Update parameters by maximizing expected complete-data log-likelihood plus log-prior:

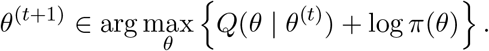

These steps are iterated until convergence.

### Extension to multi-level SLDS

For a hierarchical (multi-level) SLDS, each subject *i* has its own SLDS parameters *θ*_*i*_, latent states (*x*_*i*_, *z*_*i*_), and observations *y*_*i*_. The group SLDS parameters *η* govern the prior *π*(*θ*_*i*_ | *η*) for each subject.

To extend the EM step to the multi-level case, consider the expected complete-data log-likelihood for subject *i*:

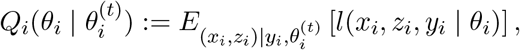

where 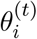 is the current parameter estimate for subject *i*. The group-level Q-function is the sum of individual ones

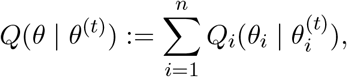

where *θ* := (*θ*, · · ·, *θ*), *θ*^(*t*)^ := (*θ*, · · ·, *θ*).

The multi-level EM proceeds in the *t*-th iteration as follows:

- **multi-level E-step**: For each subject *i* = 1, · · ·, *n* independently, compute individual Q-function 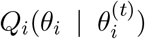 (with variational-Laplace approximation carried out individually) using current estimates 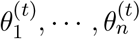. This gives expected complete-data log-likelihood per subject. The group-level *Q* function *Q*(*θ* | *θ*^(*t*)^) is then updated by summing up individual *Q*_*i*_’s.
- **multi-level M-step**: Update all subject parameters *θ* and the group parameter *η* jointly by maximizing expected complete-data log-posterior

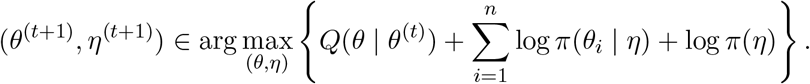

By iterating these steps, the algorithm finds each subject’s SLDS parameters and group-level parameters that (locally) maximize the hierarchical posterior. This results in a principled fit of a multi-level SLDS model, capturing both within-subject dynamics and across-subject variability.

## Appendix B: State transition measured from fMRI activity

Transition *sharpness* is a measure obtained from the model’s latent state: the flow field is estimated from the inferred latent trajectory, and the cosine between neighboring positions. Here, we asked the same question based on ROI time series (i.e., fMRI 254-dimensional space), taking from the model only the location of the boundary.

### Main idea

At each moment the whole-brain pattern of activity is moving in some direction. Over one time step the pattern changes, and the change can be written as a vector with one entry per region. That vector is a direction of travel through the space of whole-brain patterns. The question of the main text, namely, does the flow turn round at a transition, can then be evaluated based on the data. Take the average direction of travel over the few frames before the boundary, take it again over the few frames after, and measure the angle between them. If the two directions are similar, the brain kept moving the way it had been moving, and the boundary was a smooth continuation. If they differ substantially, the brain changed course. Critically, the fitted equations of motion do not enter this computation; the model is used only to say where the boundary falls.

### Activity transition

For a run’s fMRI activity *y*_1_, …, *y*_*T*_ with *y*_*t*_ *∈* ℝ^254^, let *τ* be the first frame carrying the new state label, and average the one-step displacement over the three frames on either side of the boundary:

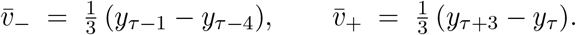

The measure is the angle between them:

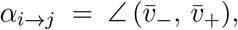

where a larger angle indicates a larger change of course. We use the angle, and not the size of the change 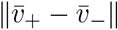, because the angle corresponds to the measure of the main text. However, for completeness we report both below.

### Reference floor value

Two directions in a 254-dimensional space (i.e., high-dimensional space) are close to perpendicular whatever the brain is doing (Vershynin, 2018). So, here, the absolute value of *α* is less informative. But a meaningful measure is how much larger the angle is than the value the same measurement determines when nothing is happening, that is, away from a state boundary. We obtained that reference from the data. For every dwell lasting at least 19 frames we took its midpoint as a pseudo-boundary, such that both windows fell inside a single state and no state change occurred between them. The reference floor for a transition was defined as the mean of the values for the two states it joins. We called this value *γ*_*i→j*_. Below, every comparison is between a transition and its own reference floor.

### the excess angle

The quantity we considered was the difference between the two,

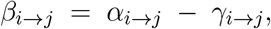

which we call the *excess angle*. It is the amount by which the brain changes course at a transition over and above the amount it appears to change course when nothing is happening.

### Averaging and inference

We enumerated every instance of the ten state transitions, discarding those lying too near the start or end of a run for the surrounding window to fit (a margin of eight frames). The angle was averaged over instances within a participant and state transition and then across the 500 participants. Confidence intervals were bootstrap intervals over participants (*B* = 10,000). The comparison below uses the same eight transitions as Fig. 5. The correspondence with the main-text measure was assessed by an exact permutation over all 8! relabelings of the transitions (rather than by a parametric *p*-value).

### fMRI activity reproduced the sharpness ordering

All transition angles differed significantly from zero, with the exception of State 1 ↦ State 5, which is the transition the main text finds smoothest by every measure. The rank correlation between the excess angle and transition sharpness, the cosine at state boundaries, was *ρ* = −0.98 across the eight transitions (Fig. S7C; exact *p* = 0.0004). Note that the relationship is negative because the two measures run in opposite directions (a more negative cosine and a larger angle both meaning a sharper transition). We also asked whether the relationship was driven by a subset of the participants. Computed separately within each participant, the same rank correlation averaged −0.54 and was negative in 94% of the 500 participants (Fig. S7D).

The size of the change, 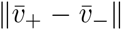, mixes the change of course with the change of speed. Measured as the angle was, as an excess over the same mid-dwell floor and across the same eight transitions, it also tracks transition sharpness (*ρ* = −0.95, exact *p* = −0.001). Direction alone therefore does a little better than direction and speed together ( −0.98 against −0.95), which is what one expects if the correspondence is carried by the change of course, but the margin is small and either quantity recovers the ordering.

**Figure S7:**
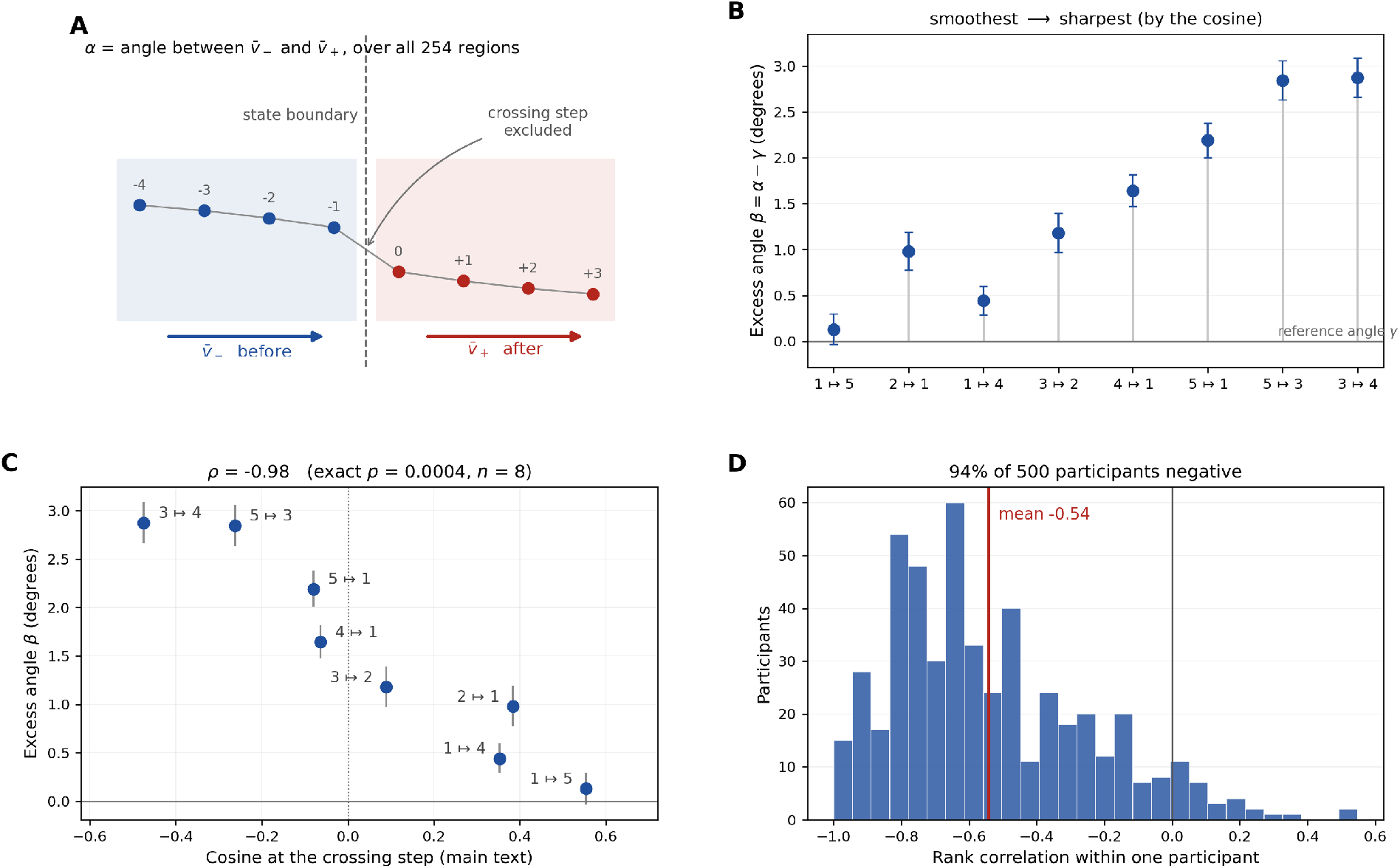
The transition measured in the regional time series. **A**. The direction in which the whole-brain pattern is travelling is averaged over the three frames before the boundary 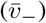 and the three frames after it 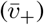, on all 254 regional time series; the step that crosses the boundary is excluded from both. The measure is based on the angle *α* between the two directions. **B**. The excess angle *β* = *α* − *γ* for each transition, with 95% bootstrap intervals over participants. The reference angle *γ* is the same measurement made at the midpoints of long dwells, where no transition occurs; it sits at zero here, and each stem shows how far that transition rises above it. Transitions are ordered by the cosine of Fig. 5, smoothest on the left. **C**. The same quantity plotted against the cosine at the crossing step. The two orderings are the reverse of each other but for one adjacent exchange, *ρ* = −0.98; the sign is negative because a more negative cosine and a larger angle both indicate a sharper transition. **D**. The same rank correlation computed separately within each participant.

## Appendix C: How far the reversal of the flow generalizes

A key result of our study was that transition sharpness varied considerably from relatively smooth to toggle-like, which was studied with six states (*K* = 6).

### The number of states

Here, we tested how robust these findings are with respect to the number of states of the model: *K* = 4 and *K* = 8. In each case, we performed a complete refit of the model (the only difference being the number of states). Fig. S8 shows the results. For the original choice of *K* = 6, four of eight transitions exhibited a reorientation of the flow in a single time step, what we called toggle-like transitions. Additional analyses with *K* = 8 states detected multiple toggle-like transitions; indeed, six of fourteen transitions. In the case of *K* = 4 states, none of the three transitions reoriented the vector field. However, the transitions exhibited a range of sharpness values, one of which was very close to reversal. The results suggest that a four-state model describes the dynamics too coarsely for a reorientation of the size present in the data to carry the cosine below zero, while models with six and eight states resolve boundaries at which it does. Taken together, these results show that the reorganization of the dynamics flow at state boundaries uncovered for *K* = 6 was not strongly dependent on the choice of number of states.

### Global signal regression

As described in Methods, we applied global signal regression during preprocessing, which is known to alter the correlation structure (Saad et al., 2012; Chai et al., 2012). Here, we refitted the model at *K* = 6 to the same 500 participants (with all parameters the same) with time series data without global signal removal. Again, we detected transition sharpness of varying steepness, including two that were toggle-like (Fig. S9). These results demonstrate that the presence of sharp transitions is not solely driven by the preprocessing choice of removing the global signal.

### Other

Transitions involving a model’s rarest state are set aside throughout because a state whose mean dwell is under three seconds does supply the eleven consecutive within-state frames the mid-dwell floor requires. The eight-state model also tests the directional claim of the Discussion. Specifically, that the brain drifts out of the dominant regime but is switched into it. In fact, neither exit from the dominant state reverses, at +0.39 and +0.35, while one of the four entries does, at −0.23.

It is tempting to assume that smooth transitions are the ones the baseline state takes part in. However, at *K* = 6 it fails in both directions at once. State 5 ↦ State 1 involves the baseline state and yet reverses, at −0.08, while State 3 ↦ State 2 does not involve it and yet does not reverse, sitting at +0.09. Overall, what appears to predict sharpness is not whether the baseline state takes part in a transition, but which way the system is traveling with respect to it.

**Figure S8:**
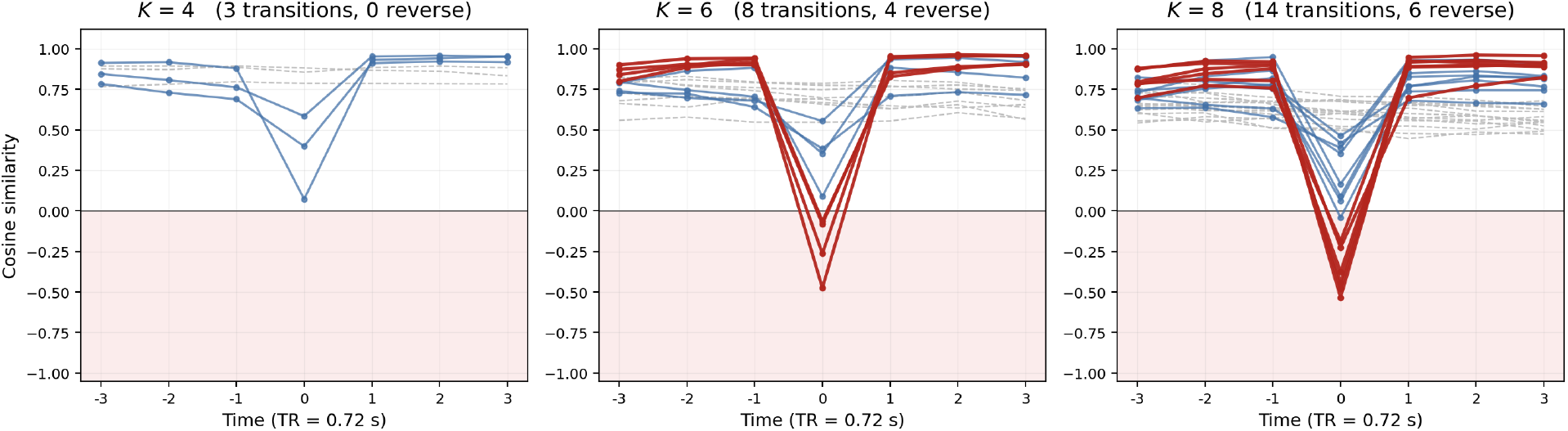
The reorientation of the flow at four, six and eight states. Each panel shows one model’s cosine between the flow field at time *t* and the field one step earlier, computed in the full ten-dimensional latent space exactly as in Fig. 5, with one curve per state transition; *t* = 0 is the boundary-crossing step. Transitions involving each model’s rarest state are excluded, leaving three, eight and fourteen. Red marks a transition whose 95% interval at the crossing step lies entirely below zero, blue one where a reversal is not established, and the shaded region below zero is where the flow has reversed. Dashed gray curves are the transitions’ own count-matched mid-dwell floors. State numbering is occupancy rank within each model and does not correspond across models.

**Figure S9:**
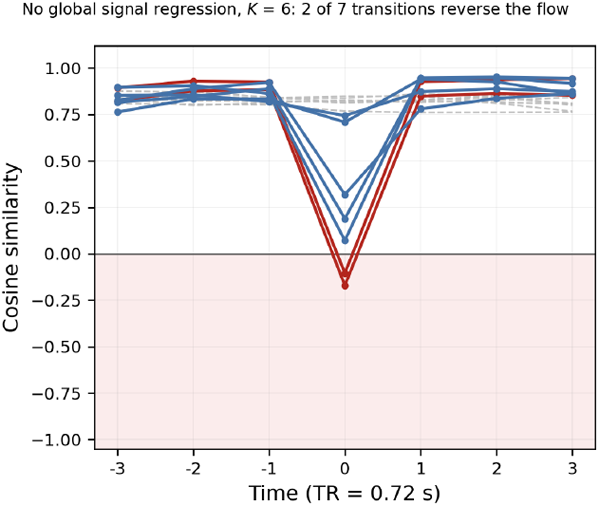
The flow reverses without global signal regression. A model fitted at *K* = 6 and *D* = 10 to the same 500 participants, the same 2,000 runs and the same 2,397,961 time points as the main text, from regional time series in which the global signal was never removed. The panel shows the cosine between the flow field at time *t* and the field one step earlier, computed in the full ten-dimensional latent space exactly as in Fig. 5, with one curve per transition; *t* = 0 is the boundary-crossing step. Red marks a transition whose 95% interval at the crossing step lies entirely below zero, blue one where a reversal is not established, and the shaded region below zero is where the flow has reversed. Two of the seven transitions reverse. Dashed gray curves are the transitions’ own count-matched mid-dwell floors. Transitions involving the model’s rarest state are excluded; that state occupies 2.7% of the time points with a mean dwell of 2.75 s, and three of the ten significant transitions involve it, none of which reverses. The curves are not labeled by state pair: this is an independent refit, and its state numbering bears no relation to the numbering used in the main text.

## Appendix D: Hemodynamics does not account for transition sharpness

BOLD does not follow neural activity instantly but is delayed and smeared in time. Suppose that every neural transition were an instantaneous jump between activity patterns, with no gradual switches at all. Early in a state’s life, the measured signal would still be climbing toward that state’s pattern; late in a long stay, the climb would have finished. Thus, apparent sharpness would be set by how far the response to the departing state’s own onset had run its course at the moment of exit. A transition leaving a state occupied only briefly would reverse the measured flow for a trivial reason, because the signal was still traveling toward the old state’s pattern when the next state change sent it toward the new one; likewise, a transition leaving the long-dwelling State 1 would look smooth for an equally simple reason, because the climb had finished and there was nothing left to reverse.

Figure S10A illustrates the account for a single, representative brain region: the pulse is the region’s activity during a stay in a state, so the duration of the pulse is the dwell and its offset is the instantaneous switch, and the smooth curve is the BOLD measured signal. A state is a pattern of activity across all regions, and at a switch every region steps toward its level in the new pattern, each with a measured signal of this shape. When those signals are still climbing at the switch, they turn around together (the pattern seen in the network time courses of Fig. 3A) and the joint trajectory in latent space doubles back (which the analysis evaluates as a reversal).

The dwell times in our data make this concern concrete. Between 57% and 72% of the dwells of State 3, State 4, and State 5 end within 5.0 s, before the response has peaked, whereas 54% of State 1’s dwells last 10.8 s or more, past its positive lobe. Sharp transitions leave short-dwelling states and smooth ones leave the long-dwelling state, which is the pattern described above. The argument of the main text, that sharp and smooth transitions share destination states and networks, rules out regional differences in response shape but does not rule out a hemodynamic origin for the state transition patterns.

The hemodynamic account comes with two implications. First, it is symmetric for a pair of states. To see why, consider the pair State 1 and State 4, and suppose the brain alternates between them by instantaneous jumps. During a stay in State 1, the measured signal travels along the line joining the two activity patterns, away from State 4’s pattern and toward State 1’s; the part of this journey still unfinished at the next jump is the “residual” response. If the jump back to State 4 arrives while the residual is large, the measured flow turns around on that same line: it was moving toward State 1 and now moves toward State 4, and the transition is scored as a reversal. A stay in State 4 is the mirror image: the signal travels the same line the other way, and an early jump back to State 1 turns it around just as fully. The two directions of a pair join the same two patterns through the same sluggish response, so, at matched dwell, they must reverse together and to the same degree (Fig. S10B). Second, it makes sharpness a function of the departing dwell time. Taken together, the implications make three predictions. Within a transition, sharpness should track the length of the departing dwell. Across transitions, the sharp-versus-smooth difference should vanish once dwell is matched. And no reversal should survive when the departing state was occupied long enough for the measured signal to finish climbing.

We tested both implications by transition sharpness (Fig. 5) after splitting each transition’s instances by the length of the departing dwell (called “stratum” next): seven frames or fewer (at most 5.0 s), eight to fourteen frames, and fifteen frames or more (at least 10.8 s). Everything else was left exactly as in Fig. 5: each stratum was compared against its own count-matched mid-dwell floor, and intervals came from the same participant subsampling. Instances whose departing dwell began at the first frame of a run were excluded, because their true onset is unknown; they are under one percent of instances. Two cells of the design cannot be estimated and are marked as such in Table S3: dwells of fifteen frames or more are rare in State 5 (4% of its dwells), so the long strata of State 5 ↦ State 1 and State 5 ↦ State 3 retain too few instances.

**Figure S10:**
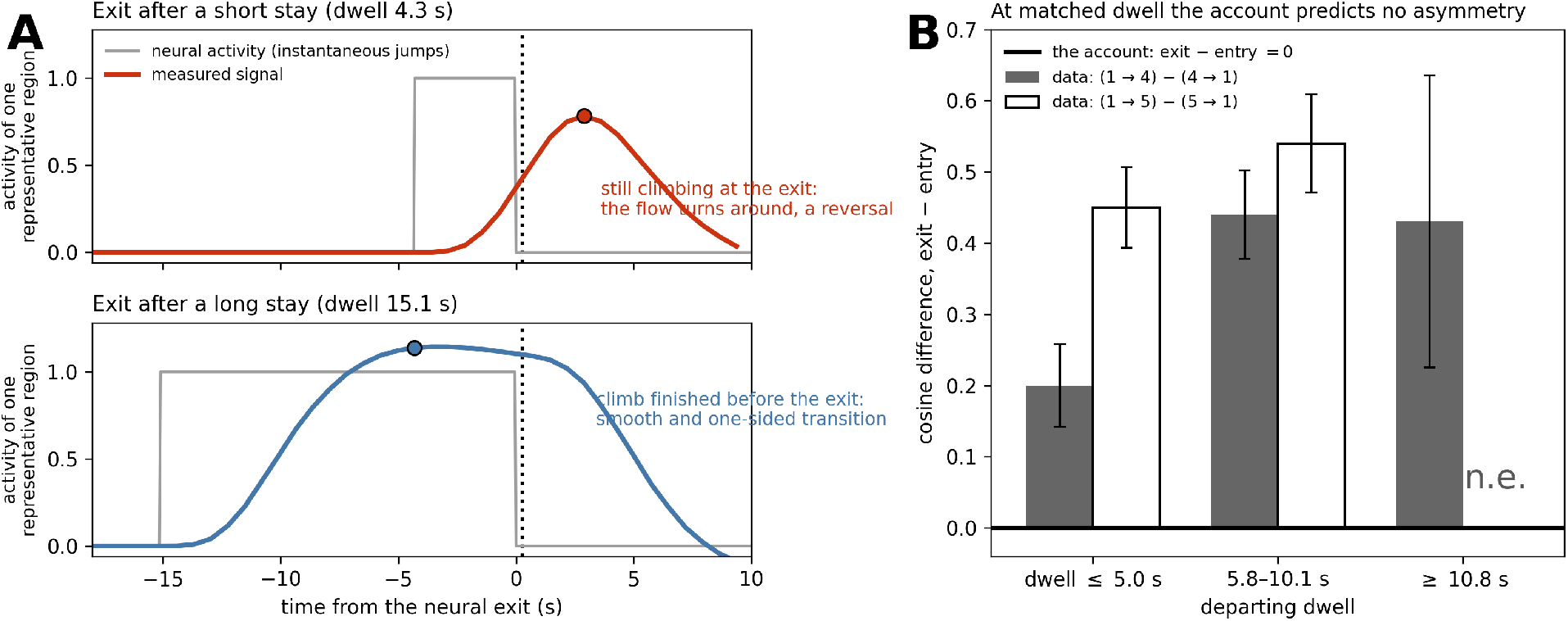
Hemodynamic potential account and its test. **A**. The account for a single, representative brain region. The pulse (gray) is the region’s neural activity during one stay in a state: the duration of the pulse is the dwell, and its offset is the instantaneous switch to the next state. The curve (color) is the measured signal, sampled at the repetition time of 0.72 s. After a short stay (top), the signal is still climbing when the switch arrives, so the measured flow turns around at the boundary the analysis detects (dot) and the transition is scored as a reversal; after a long stay (bottom), the climb has finished well before the exit (dot) and the measured profile is smooth and one-sided. When multiple regions traces a curve of this shape between its levels in the two state patterns, the reversal in the panel appears in latent space as the trajectory doubling back (i.e., reversal of the vector fields). **B**. The hemodynamic account’s decisive implication: at matched departing dwell, the exit and the entry of a pair must be equally sharp, so their difference should be zero at every dwell (the zero line). The measured differences (bars; 95% intervals)lie far above zero in every stratum; the long stratum of the State 1–State 5 pair is not estimable (n.e.).

The test comes out in favor of the main text (Table S3). With departing dwell matched, the asymmetry between the two directions of a pair survives in every stratum where it can be measured. Consider the short stratum, in which every departing state was occupied for at most 5.0 s, so the hemodynamic account predicts a reversal for both directions of every pair. The exits from State 1 stay positive while the entries into it reverse, and the difference between the two directions, which the account cannot produce at all, is +0.20 for the State 1–State 4 pair and +0.45 for State 1–State 5 (*z* = 6.8 and 15.5). At medium dwell the two differences are +0.44 and +0.54, equally far from zero. In the long stratum only the State 1–State 4 pair can be formed, because dwells of fifteen frames or more are rare in State 5; that single comparison gives +0.43 (*z* = 4.1), the same answer as the other two strata. The long stratum also shows the complementary failure: State 3 ↦State 4 still reverses after departing dwells of 10.8 s or more, where the hemodynamic account predicts a smooth exit. Both implications fail together, and the direction-dependence reported in the main text is therefore a property of the dynamics rather than of the measurement.

At the same time, some hemodynamic contribution cannot be excluded, and Table S3 shows where it plausibly acts: the exits from State 1 grow smoother as the departing dwell lengthens, from +0.18 to +0.38 for State 1 ↦ State 4 and from +0.37 to +0.58 for State 1 ↦ State 5, which is what a delayed and smeared response should add. What the account does not do is produce our results: it does not set the sign of the State 1 exits, it does not remove the reversal of State 3 ↦ State 4 after long dwells, and it cannot, by construction, tell the two directions of a pair apart. We conclude that the sharp-versus-smooth split, and in particular its dependence on direction, is a property of the underlying dynamics rather than of the measurement.

**Table S3:**
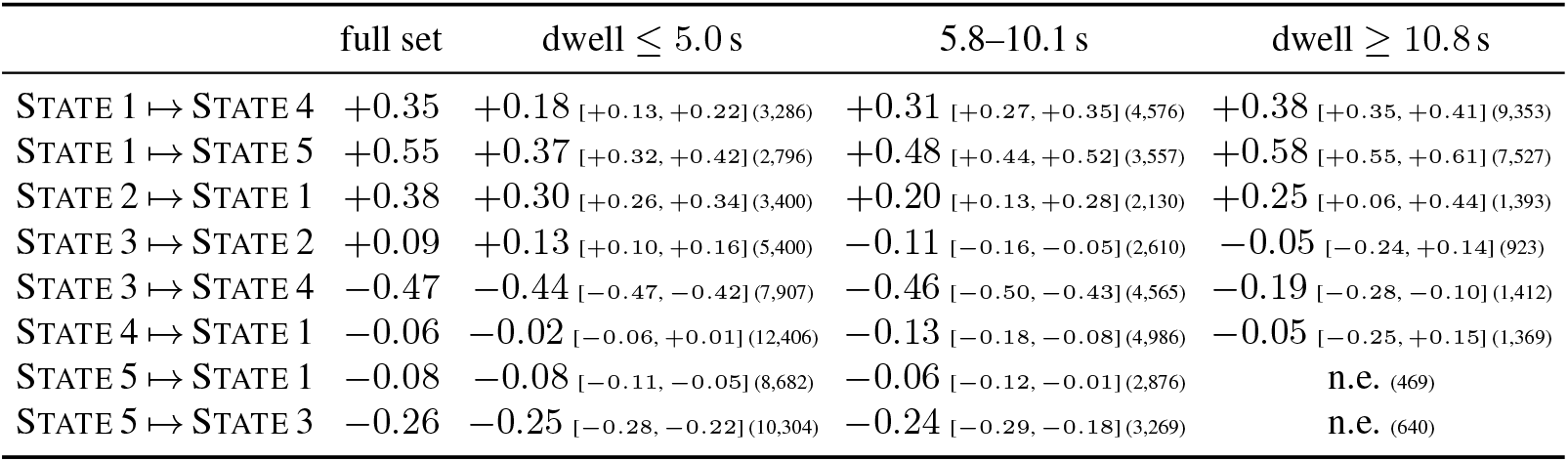
Transition sharpness for different lengths of the original state departing dwell, with 95% intervals and instance counts. The strata are seven frames or fewer, eight to fourteen, and fifteen or more, at the repetition time of 0.72 s; the full-set column repeats Fig. 5, whose intervals are given there. Exits from State 1 stay positive in every stratum, including the short one, where the hemodynamic account predicts reversals as deep as the entries’; State 3 ↦ State 4 still reverses in the long stratum. Cells marked n.e. (not estimable) retain too few instances because dwells of fifteen frames or more are rare in State 5.

## References

Elena A. Allen, Eswar Damaraju, Sergey M. Plis, Erik B. Erhardt, Tom Eichele, and Vince D. Calhoun. Tracking whole-brain connectivity dynamics in the resting state. Cerebral Cortex, 24(3):663–676, 2012. doi: 10.1093/cercor/bhs352.

Jeffrey Alstott, Michael Breakspear, Patric Hagmann, Leila Cammoun, and Olaf Sporns. Modeling the impact of lesions in the human brain. PLoS Computational Biology, 5(6):e1000408, 2009. doi: 10.1371/journal.pcbi.1000408.

Adam P. Baker, Matthew J. Brookes, Iead A. Rezek, Stephen M. Smith, Timothy Behrens, Penny J. Probert Smith, and Mark Woolrich. Fast transient networks in spontaneous human brain activity. eLife, 3:e01867, 2014. doi: 10.7554/eLife.01867.

David Barber. Expectation correction for smoothed inference in switching linear dynamical systems. Journal of Machine Learning Research, 7(89):2515–2540, 2006.

Yoav Benjamini and Yosef Hochberg. Controlling the false discovery rate: A practical and powerful approach to multiple testing. Journal of the Royal Statistical Society Series B (Methodological), 57(1): 289–300, 1995.

Taylor Bolt, Jason S. Nomi, Danilo Bzdok, Jorge A. Salas, Catie Chang, B. T. Thomas Yeo, Lucina Q. Uddin, and Shella D. Keilholz. A parsimonious description of global functional brain organization in three spatiotemporal patterns. Nature Neuroscience, 25(8):1093–1103, 2022. doi: 10.1038/s41593-022-01118-1.

Joana Cabral, Morten L. Kringelbach, and Gustavo Deco. Exploring the network dynamics underlying brain activity during rest. Progress in Neurobiology, 114:102–131, 2014.

Xiaoqian J. Chai, Alfonso N. Castañón, Dost Öngür, and Susan Whitfield-Gabrieli. Anticorrelations in resting state networks without global signal regression. NeuroImage, 59(2):1420–1428, 2012.

Kaixin Chen, Chuanjun Li, Wei Sun, Yu Tao, Ruilin Wang, Wei Hou, and Dong-Qiang Liu. Hidden Markov modeling reveals prolonged “baseline” state and shortened antagonistic state across the adult lifespan. Cerebral Cortex, 32(2):439–453, 2022.

Eli J. Cornblath, Arian Ashourvan, Jason Z. Kim, Richard F. Betzel, Rastko Ciric, Azeez Adebimpe, Gra-ham L. Baum, Xiaosong He, Kosha Ruparel, Tyler M. Moore, Ruben C. Gur, Raquel E. Gur, Russell T. Shinohara, David R. Roalf, Theodore D. Satterthwaite, and Danielle S. Bassett. Temporal sequences of brain activity at rest are constrained by white matter structure and modulated by cognitive demands. Communications Biology, 3:261, 2020. doi: 10.1038/s42003-020-0961-x.

Robert W. Cox. AFNI: software for analysis and visualization of functional magnetic resonance neuroimages. Computers and Biomedical Research, 29(3):162–173, 1996.

Eswar Damaraju, Elena A. Allen, Aysenil Belger, Judith M. Ford, Sarah McEwen, Daniel H. Mathalon, Bryon A. Mueller, Godfrey D. Pearlson, Steven G. Potkin, Adrian Preda, Jessica A. Turner, Jatin G. Vaidya, Theo G. van Erp, and Vince D. Calhoun. Dynamic functional connectivity analysis reveals transient states of dysconnectivity in schizophrenia. NeuroImage: Clinical, 5:298–308, 2014. doi: 10.1016/j.nicl.2014.07.003.

Gustavo Deco and Viktor K. Jirsa. Ongoing cortical activity at rest: Criticality, multistability, and ghost attractors. Journal of Neuroscience, 32(10):3366–3375, 2012.

Gustavo Deco, Josephine Cruzat, Joana Cabral, Enzo Tagliazucchi, Helmut Laufs, Nikos K. Logothetis, and Morten L. Kringelbach. Awakening: Predicting external stimulation to force transitions between different brain states. Proceedings of the National Academy of Sciences, 116(36):18088–18097, 2019. doi: 10.1073/pnas.1905534116.

Stanislas Dehaene and Jean-Pierre Changeux. Experimental and theoretical approaches to conscious processing. Neuron, 70(2):200–227, 2011. doi: 10.1016/j.neuron.2011.03.018.

Arthur P. Dempster, Nan M. Laird, and Donald B. Rubin. Maximum likelihood from incomplete data via the EM algorithm. Journal of the Royal Statistical Society Series B (Methodological), 39(1):1–38, 1977.

Takahiro Ezaki, Michiko Sakaki, Takamitsu Watanabe, and Naoki Masuda. Age-related changes in the ease of dynamical transitions in human brain activity. Human Brain Mapping, 39(6):2673–2688, 2018. doi: 10.1002/hbm.24033.

Michael D. Fox, Abraham Z. Snyder, Justin L. Vincent, Maurizio Corbetta, David C. Van Essen, and Marcus E. Raichle. The human brain is intrinsically organized into dynamic, anticorrelated functional networks. Proceedings of the National Academy of Sciences, 102(27):9673–9678, 2005. doi: 10.1073/pnas.0504136102.

Zoubin Ghahramani and Geoffrey E. Hinton. Variational learning for switching state-space models. Neural Computation, 12(4):831–864, 2000. doi: 10.1162/089976600300015619.

Abigail S. Greene, Corey Horien, Daniel Barson, Dustin Scheinost, and R. Todd Constable. Why is everyone talking about brain state? Trends in Neurosciences, 46(7):508–524, 2023. doi: 10.1016/j.tins.2023.04.001.

Shi Gu, Fabio Pasqualetti, Matthew Cieslak, Qawi K. Telesford, Alfred B. Yu, Ari E. Kahn, John D. Medaglia, Jean M. Vettel, Michael B. Miller, Scott T. Grafton, and Danielle S. Bassett. Controllability of structural brain networks. Nature Communications, 6:8414, 2015. doi: 10.1038/ncomms9414.

Shi Gu, Richard F. Betzel, Marcelo G. Mattar, Matthew Cieslak, Philip R. Delio, Scott T. Grafton, Fabio Pasqualetti, and Danielle S. Bassett. Optimal trajectories of brain state transitions. NeuroImage, 148: 305–317, 2017. doi: 10.1016/j.neuroimage.2017.01.003.

Daniel Gutierrez-Barragan, Julian S. B. Ramirez, Stefano Panzeri, Ting Xu, and Alessandro Gozzi. Evolutionarily conserved fMRI network dynamics in the mouse, macaque, and human. Nature Communications, 15(1):8518, 2024. doi: 10.1038/s41467-024-52721-8.

Daniel A. Handwerker, John M. Ollinger, and Mark D’Esposito. Variation of BOLD hemodynamic responses across subjects and brain regions and their effects on statistical analyses. NeuroImage, 21(4): 1639–1651, 2004. doi: 10.1016/j.neuroimage.2003.11.029.

R. Matthew Hutchison, Thilo Womelsdorf, Elena A. Allen, Peter A. Bandettini, Vince D. Calhoun, Maur-izio Corbetta, Stefania Della Penna, Jeff H. Duyn, Gary H. Glover, Javier Gonzalez-Castillo, Daniel A. Handwerker, Shella Keilholz, Vesa Kiviniemi, David A. Leopold, Francesco de Pasquale, Olaf Sporns, Martin Walter, and Catie Chang. Dynamic functional connectivity: Promise, issues, and interpretations. NeuroImage, 80:360–378, 2013. doi: 10.1016/j.neuroimage.2013.05.079.

Lauren M. Jones, Alfredo Fontanini, Brian F. Sadacca, Paul Miller, and Donald B. Katz. Natural stimuli evoke dynamic sequences of states in sensory cortical ensembles. Proceedings of the National Academy of Sciences, 104(47):18772–18777, 2007. doi: 10.1073/pnas.0705546104.

Fikret I. Karahanoğlu and Dimitri Van De Ville. Transient brain activity disentangles fMRI resting-state dynamics in terms of spatially and temporally overlapping networks. Nature Communications, 6:7751, 2015. doi: 10.1038/ncomms8751.

Matthew T. Kaufman, Mark M. Churchland, Stephen I. Ryu, and Krishna V. Shenoy. Cortical activity in the null space: permitting preparation without movement. Nature Neuroscience, 17(3):440–448, 2014. doi: 10.1038/nn.3643.

Shella D. Keilholz. The neural basis of time-varying resting-state functional connectivity. Brain Connectivity, 4(10):769–779, 2014.

Shella D. Keilholz, Matthew E. Magnuson, Wen-Ju Pan, Millicent Willis, and Garth J. Thompson. Dynamic properties of functional connectivity in the rodent. Brain Connectivity, 3(1):31–40, 2013.

Mikail Khona and Ila R. Fiete. Attractor and integrator networks in the brain. Nature Reviews Neuroscience, 23(12):744–766, 2022. doi: 10.1038/s41583-022-00642-0.

Giancarlo La Camera, Alfredo Fontanini, and Luca Mazzucato. Cortical computations via metastable activity. Current Opinion in Neurobiology, 58:37–45, 2019. doi: 10.1016/j.conb.2019.06.007.

Kenneth W. Latimer, Jacob L. Yates, Miriam L. R. Meister, Alexander C. Huk, and Jonathan W. Pillow. Single-trial spike trains in parietal cortex reveal discrete steps during decision-making. Science, 349 (6244):184–187, 2015. doi: 10.1126/science.aaa4056.

Kenneth W. Latimer, Jacob L. Yates, Miriam L. R. Meister, Alexander C. Huk, and Jonathan W. Pillow. Response to comment on “single-trial spike trains in parietal cortex reveal discrete steps during decisionmaking”. Science, 351(6280):1406, 2016. doi: 10.1126/science.aad3596.

Kangjoo Lee, Jie Lisa Ji, Clára Fonteneau, Lucie Berkovitch, Masih Rahmati, Lining Pan, Grega Repovš, John H. Krystal, John D. Murray, and Alan Anticevic. Human brain state dynamics are highly reproducible and associated with neural and behavioral features. PLOS Biology, 22(9):1–34, 2024. doi: 10.1371/journal.pbio.3002808.

Nora Leonardi and Dimitri Van De Ville. On spurious and real fluctuations of dynamic functional connectivity during rest. NeuroImage, 104:430–436, 2015. doi: 10.1016/j.neuroimage.2014.09.007.

Raphaël Liégeois, Jingwei Li, Ru Kong, Csaba Orban, Dimitri Van De Ville, Tian Ge, Mert R. Sabuncu, and B. T. Thomas Yeo. Resting brain dynamics at different timescales capture distinct aspects of human behavior. Nature Communications, 10(1):2317, 2019. doi: 10.1038/s41467-019-10317-7.

Scott Linderman, Matthew Johnson, Andrew Miller, Ryan Adams, David Blei, and Liam Paninski. Bayesian learning and inference in recurrent switching linear dynamical systems. In Proceedings of the 20th International Conference on Artificial Intelligence and Statistics, volume 54 of Proceedings of Machine Learning Research, pages 914–922, 2017.

Scott Linderman, Annika Nichols, David Blei, Manuel Zimmer, and Liam Paninski. Hierarchical recurrent state space models reveal discrete and continuous dynamics of neural activity in C. elegans. bioRxiv, 2019. doi: 10.1101/621540.

Xiao Liu, Nanyin Zhang, Catie Chang, and Jeff H. Duyn. Co-activation patterns in resting-state fMRI signals. NeuroImage, 180:485–494, 2018.

David G. Luenberger. Introduction to Dynamic Systems: Theory, Models, and Applications. Wiley, 1979.

Daniel J. Lurie, Daniel Kessler, Danielle S. Bassett, Richard F. Betzel, Michael Breakspear, Shella Kheil-holz, Aaron Kucyi, Raphaël Liégeois, Martin A. Lindquist, Anthony R. McIntosh, Russell A. Poldrack, James M. Shine, William H. Thompson, Natalia Z. Bielczyk, Linda Douw, Dominik Kraft, Robyn L. Miller, Muthuraman Muthuraman, Lorenzo Pasquini, Adeel Razi, and et al. Questions and controversies in the study of time-varying functional connectivity in resting fMRI. Network Neuroscience, 4(1):30–69, 2020.

Waqas Majeed, Matthew Magnuson, Wendy Hasenkamp, Hillary Schwarb, Eric H. Schumacher, Lawrence Barsalou, and Shella D. Keilholz. Spatiotemporal dynamics of low frequency BOLD fluctuations in rats and humans. NeuroImage, 54(2):1140–1150, 2011.

Eric Maltbie, Behnaz Yousefi, Xiaodi Zhang, Amrit Kashyap, and Shella Keilholz. Comparison of resting-state functional MRI methods for characterizing brain dynamics. Frontiers in Neural Circuits, 16:681544, 2022.

Luca Mazzucato, Alfredo Fontanini, and Giancarlo La Camera. Dynamics of multistable states during ongoing and evoked cortical activity. Journal of Neuroscience, 35(21):8214–8231, 2015. doi: 10.1523/JNEUROSCI.4819-14.2015.

Paul Miller. Itinerancy between attractor states in neural systems. Current Opinion in Neurobiology, 40: 14–22, 2016.

Joyneel Misra and Luiz Pessoa. Brain dynamics and spatiotemporal trajectories during threat processing. eLife, 14:RP102539, 2025. doi: 10.7554/eLife.102539.2.

Sang Min Oh, James M. Rehg, Tucker Balch, and Frank Dellaert. Learning and inferring motion patterns using parametric segmental switching linear dynamic systems. International Journal of Computer Vision, 77(1):103–124, 2008. doi: 10.1007/s11263-007-0062-z.

Xiaowei Peng, Quanying Liu, Catie S. Hubbard, Danhong Wang, Wanwan Zhu, Michael D. Fox, and Hes-heng Liu. Robust dynamic brain coactivation states estimated in individuals. Science Advances, 9(3): eabq8566, 2023. doi: 10.1126/sciadv.abq8566.

Mikhail I. Rabinovich, Pablo Varona, Allen I. Selverston, and Henry D. I. Abarbanel. Dynamical principles in neuroscience. Reviews of Modern Physics, 78(4):1213–1265, 2006. doi: 10.1103/RevModPhys.78.1213.

Ryan V. Raut, Abraham Z. Snyder, Anish Mitra, Dov Yellin, Naotaka Fujii, Rafael Malach, and Marcus E. Raichle. Global waves synchronize the brain’s functional systems with fluctuating arousal. Science Advances, 7(30):eabf2709, 2021. doi: 10.1126/sciadv.abf2709.

Stefano Recanatesi, Ulises Pereira-Obilinovic, Masayoshi Murakami, Zachary Mainen, and Luca Mazzucato. Metastable attractors explain the variable timing of stable behavioral action sequences. Neuron, 110 (1):139–153, 2022. doi: 10.1016/j.neuron.2021.10.011.

Ziad S. Saad, Stephen J. Gotts, Kevin Murphy, Gang Chen, Hang Joon Jo, Alex Martin, and Robert W. Cox. Trouble at rest: how correlation patterns and group differences become distorted after global signal regression. Brain Connectivity, 2(1):25–32, 2012.

Maria V. Sanchez-Vives and David A. McCormick. Cellular and network mechanisms of rhythmic recurrent activity in neocortex. Nature Neuroscience, 3(10):1027–1034, 2000.

Clifford B. Saper, Patrick M. Fuller, Nigel P. Pedersen, Jun Lu, and Thomas E. Scammell. Sleep state switching. Neuron, 68(6):1023–1042, 2010. doi: 10.1016/j.neuron.2010.11.032.

Alexander Schaefer, Ru Kong, Evan M. Gordon, Timothy O. Laumann, Xi-Nian Zuo, Avram J. Holmes, Simon B. Eickhoff, and B. T. Thomas Yeo. Local-global parcellation of the human cerebral cortex from intrinsic functional connectivity MRI. Cerebral Cortex, 28(9):3095–3114, 2017. doi: 10.1093/cercor/bhx179.

Marten Scheffer, Jordi Bascompte, William A. Brock, Victor Brovkin, Stephen R. Carpenter, Vasilis Dakos, Hermann Held, Egbert H. van Nes, Max Rietkerk, and George Sugihara. Early-warning signals for critical transitions. Nature, 461(7260):53–59, 2009. doi: 10.1038/nature08227.

João D. Semedo, Amin Zandvakili, Christian K. Machens, Byron M. Yu, and Adam Kohn. Cortical areas interact through a communication subspace. Neuron, 102(1):249–259, 2019. doi: 10.1016/j.neuron.2019.01.026.

Michael N. Shadlen, Roozbeh Kiani, William T. Newsome, Joshua I. Gold, Daniel M. Wolpert, Ariel Zylber-berg, Jochen Ditterich, Victor de Lafuente, Tianming Yang, and Jamie Roitman. Comment on “single-trial spike trains in parietal cortex reveal discrete steps during decision-making”. Science, 351(6280):1406, 2016. doi: 10.1126/science.aad3242.

Heather Shappell, Brian S. Caffo, James J. Pekar, and Martin A. Lindquist. Improved state change estimation in dynamic functional connectivity using hidden semi-Markov models. NeuroImage, 191:243–257, 2019. doi: 10.1016/j.neuroimage.2019.02.013.

Jason F. Smith, Ajay Pillai, Kewei Chen, and Barry Horwitz. Identification and validation of effective connectivity networks in functional magnetic resonance imaging using switching linear dynamic systems. NeuroImage, 52(3):1027–1040, 2010.

Stephen M. Smith, Christian F. Beckmann, Jesper Andersson, Edward J. Auerbach, Janine Bijsterbosch, Gwenaëlle Douaud, Eugene Duff, David A. Feinberg, Ludovica Griffanti, Michael P. Harms, Michael Kelly, Timothy Laumann, Karla L. Miller, Steen Moeller, Steve Petersen, Jonathan Power, Gholamreza Salimi-Khorshidi, Abraham Z. Snyder, An T. Vu, Mark W. Woolrich, and et al. Resting-state fMRI in the Human Connectome Project. NeuroImage, 80:144–168, 2013.

Hayoung Song, Won Mok Shim, and Monica D. Rosenberg. Large-scale neural dynamics in a shared low-dimensional state space reflect cognitive and attentional dynamics. eLife, 12:e85487, 2023. doi: 10.7554/eLife.85487.

Jalil Taghia, Weidong Cai, Srikanth Ryali, John Kochalka, Jonathan Nicholas, Tianwen Chen, and Vinod Menon. Uncovering hidden brain state dynamics that regulate performance and decision-making during cognition. Nature Communications, 9(1):2505, 2018. doi: 10.1038/s41467-018-04723-6.

Enzo Tagliazucchi and Helmut Laufs. Decoding wakefulness levels from typical fMRI resting-state data reveals reliable drifts between wakefulness and sleep. Neuron, 82(3):695–708, 2014. doi: 10.1016/j.neuron.2014.03.020.

Ye Tian, Daniel S. Margulies, Michael Breakspear, and Andrew Zalesky. Topographic organization of the human subcortex unveiled with functional connectivity gradients. Nature Neuroscience, 23(11):1421–1432, 2020. doi: 10.1038/s41593-020-00711-6.

Emmanuelle Tognoli and J. A. Scott Kelso. The metastable brain. Neuron, 81(1):35–48, 2014.

Roman Vershynin. High-Dimensional Probability: An Introduction with Applications in Data Science. Cambridge University Press, 2018. doi: 10.1017/9781108231596.

Diego Vidaurre, Stephen M. Smith, and Mark W. Woolrich. Brain network dynamics are hierarchically organized in time. Proceedings of the National Academy of Sciences, 114(48):12827–12832, 2017. doi: 10.1073/pnas.1705120114.

Diego Vidaurre, Romesh Abeysuriya, Robert Becker, Andrew J. Quinn, Fidel Alfaro-Almagro, Stephen M. Smith, and Mark W. Woolrich. Discovering dynamic brain networks from big data in rest and task. NeuroImage, 180:646–656, 2018a.

Diego Vidaurre, Laurence T. Hunt, Andrew J. Quinn, Benjamin A. E. Hunt, Matthew J. Brookes, Anna C. Nobre, and Mark W. Woolrich. Spontaneous cortical activity transiently organises into frequency specific phase-coupling networks. Nature Communications, 9:2987, 2018b. doi: 10.1038/s41467-018-05316-z.

Nguyen Xuan Vinh, Julien Epps, and James Bailey. Information theoretic measures for clusterings comparison: Variants, properties, normalization and correction for chance. Journal of Machine Learning Research, 11(95):2837–2854, 2010.

Chenhao Wang, Ju Lynn Ong, Amiya Patanaik, Juan Zhou, and Michael W. L. Chee. Spontaneous eyelid closures link vigilance fluctuation with fMRI dynamic connectivity states. Proceedings of the National Academy of Sciences, 113(34):9653–9658, 2016. doi: 10.1073/pnas.1523980113.

Takamitsu Watanabe, Satoshi Hirose, Hiroyuki Wada, Yoshio Imai, Toru Machida, Ichiro Shirouzu, Seiki Konishi, Yasushi Miyashita, and Naoki Masuda. Energy landscapes of resting-state brain networks. Frontiers in Neuroinformatics, 8:12, 2014. doi: 10.3389/fninf.2014.00012.

B. T. Thomas Yeo, Fenna M. Krienen, Jorge Sepulcre, Mert R. Sabuncu, Danial Lashkari, Marisa Hollinshead, Joshua L. Roffman, Jordan W. Smoller, Lilla Zöllei, Jonathan R. Polimeni, Bruce Fischl, Hesheng Liu, and Randy L. Buckner. The organization of the human cerebral cortex estimated by intrinsic functional connectivity. Journal of Neurophysiology, 106(3):1125–1165, 2011.

David Zoltowski, Jonathan Pillow, and Scott Linderman. A general recurrent state space framework for modeling neural dynamics during decision-making. In Proceedings of the 37th International Conference on Machine Learning, volume 119 of Proceedings of Machine Learning Research, pages 11680–11691, 2020.

